# Human bone mesoscale 3D structure revisited by plasma focused ion beam serial sectioning

**DOI:** 10.1101/2020.07.01.180729

**Authors:** Dakota Marie Binkley, Joseph Deering, Hui Yuan, Aurélien Gourrier, Kathryn Grandfield

**Author notes:** Corresponding Author: Kathryn Grandfield, Department of Materials Science and Engineering, McMaster University, 1280 Main Street West, Hamilton, ON, L8S 4L7, Canada.

## Abstract

Visualizing bone mineralization and collagen microfibril organization at intermediate scales between the nanometer and the 100s of microns range, the mesoscale, is still an important challenge. Similarly, visualizing cellular components which locally affect the tissue structure requires a precision of a few tens of nanometers at maximum while spanning several tens of micrometers. To address this issue, we employed a plasma focused ion beam (PFIB) equipped with a scanning electron microscope (SEM) to sequentially section nanometer-scale layers of demineralized and mineralized human femoral lamellar bone over volumes of approximately 46 × 40 × 9 μm^3^, and 29 × 26 × 9 μm^3^, respectively. This large scale view retained high enough resolution to visualize the collagen microfibrils while partly visualizing the lacuno-canalicular network (LCN) in three-dimensions (3D). We showed that serial sectioning can be performed on mineralized sections, and does not require demineralization. Moreover, this method revealed ellipsoidal mineral clusters, noted by others in high resolution studies, as a ubiquitous motif in lamellar bone over tens of microns, suggesting a heterogeneous and yet regular pattern of mineral deposition past the single collagen fibril level. These findings are strong evidence for the need to revisit bone mineralization over multi-length scales.

**Graphical Abstract:** 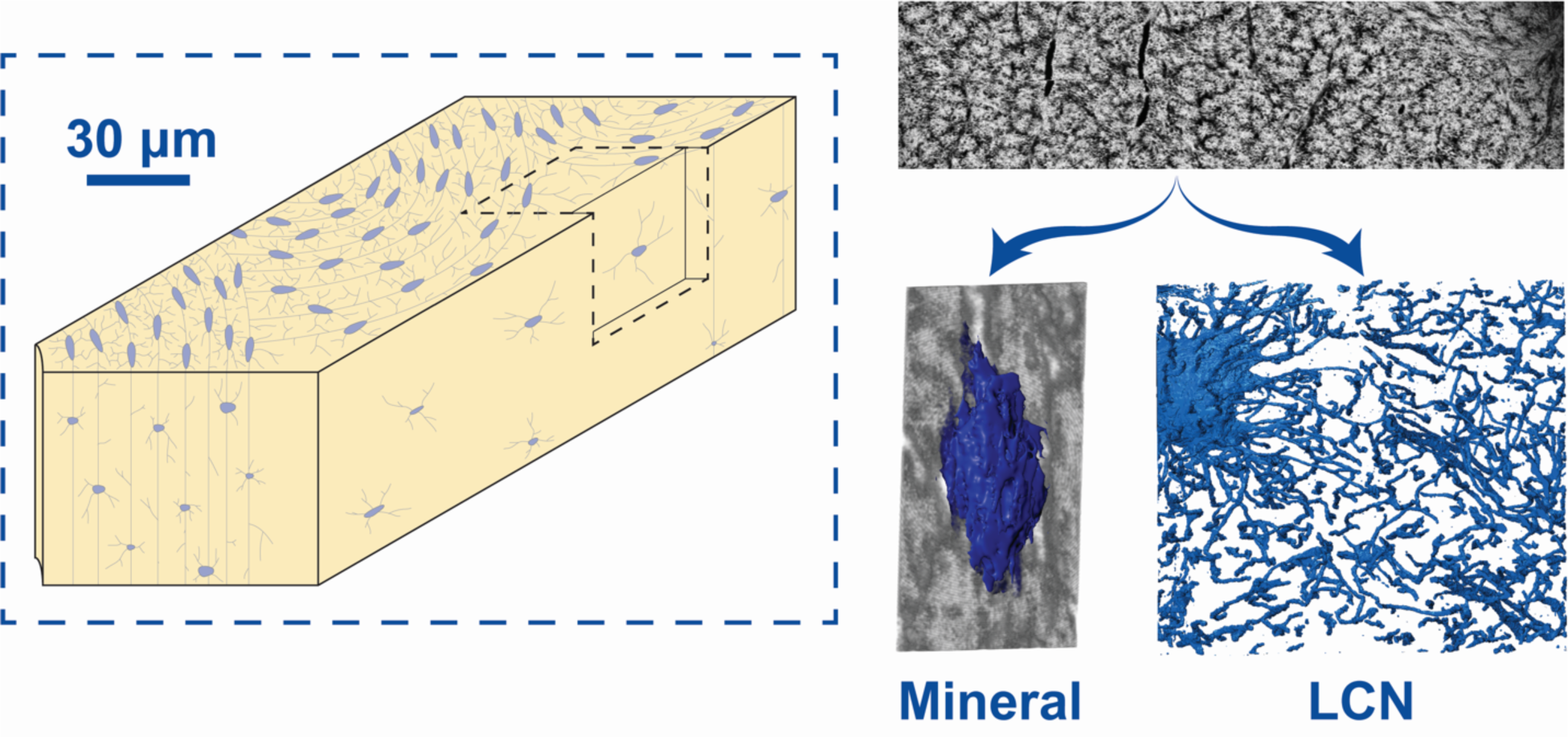

## Introduction

Bone’s remarkable mechanical properties are essentially determined by its macro-to-microarchitecture and tissue properties (Currey, 2002). Mineralized collagen microfibrils with typical diameters of 100 nm are considered to be the fundamental building block of this multiscale structural hierarchy and tissue mechanics is primarily determined by fibril orientation and mineral density (Granke et al., 2013; Wagermaier et al., 2006; Weiner et al., 1999). Thus, the precise organization of collagen microfibrils and of the mineral phase in healthy and pathological bone tissue is still an active area of research.

In addition to the intrinsic microfibril organization patterns in a 1-10 µm range, resulting from tight biological control and self-assembly mechanisms, periodic modulations can be observed over distances up to 100 µm in lamellar bone of many species. This characteristic length scale corresponds to the presence of vascular channels in human cortical bone around which the mineralized microfibrils are organized with high symmetrical regularity (Giraud-Guille, 1988; Wagermaier et al., 2006; Weiner et al., 1999). It also corresponds to the typical thickness of trabecular bone which is mostly found in the epiphyses of long bones, between the two surfaces of flat bones and also consolidates irregular bones. This high degree of organization up to the microscopic scale is, however, partly perturbed by dendritic osteocytes embedded in the tissue, whose cell bodies are usually distant by ∼ 25 µm on average, depending on the tissue symmetry (Hannah et al., 2010; Kollmannsberger et al., 2017; Repp et al., 2017). This dense, highly interconnected cellular network is a key player for bone functions (Bonewald, 2011) and potentially constitutes a source of local disorganization with respect to higher levels of symmetry. It is therefore becoming evident that analyzing the sub-microscopic structure of bone tissue also requires visualizing the lacuno-canalicular porosity network (LCN) hosting the osteocytes (Kerschnitzki et al., 2011) as well as the cellular details.

While a plethora of characterization tools exist for bone (Georgiadis et al., 2016)visualizing ultrastructural details with sufficient resolution to distinguish 100 nm diameter mineralized microfibrils while probing sufficiently large volumes with representative LCN organizations (in the 100 µm range) is still an important technical challenge.

A subset of ultrastructural characterization explores collagen-mineral arrangement with electron microscopy, chiefly transmission electron microscopy (TEM) as well as electron tomography (Grandfield et al., 2018; Landis et al., 1996; McNally et al., 2012; Reznikov et al., 2018; Weiner and Traub, 1992), which allows visualizing small volumes of up to 200 nm thick sections with sub-nanometer to angstrom resolution. Although TEM techniques have been central to defining collagen-mineral arrangement and mineral platelet morphology, even electron tomography is limited by geometrical restrictions and specimen thickness requirements, leading to missing wedge artifacts (Wang et al., 2016) which conflate reconstructions and therefore, some conclusions made via this method. Recently, another electron microscopy approach has surfaced as a highly complementary tool of TEM in bone hierarchical investigations, namely, focused ion beam (FIB) scanning electron microscopy (SEM), usually abbreviated to FIB-SEM, but also called FIB-SEM serial sectioning or FIB-SEM tomography. In this method, a heavy ion (e.g. gallium) beam is used to sequentially mill material, the block-face of which is then imaged with a coincident electron beam. FIB-SEM enables visualization of larger volumes than TEM (approximately 10 x 10 x 10 µm) which partly compensates for the loss in resolution to a few nanometers. Indeed, FIB-SEM serial-sectioning has been demonstrated on embedded or cryogenically preserved tissues including, mineralized turkey tendon (Zou et al., 2019), human, pig and rat lamellar bone (Reznikov et al., 2014b, 2014c, 2013), human trabecular bone (Reznikov et al., 2014a), mandibular bone of the minipig (Maria et al., 2019), zebrafish larvae (Akiva et al., 2019; Silvent et al., 2017), embryonic chicken bone (Kerschnitzki et al., 2016a) and the interface of cementum and periodontal ligament (Hirashima et al., 2020b, 2020a) to name a few. Nearly all studies utilize demineralized bone and were therefore focused on the collagen microfibrils. Very few studies capture large sections of the LCN, which were therefore performed with a much lower spatial resolution than technically possible, to acquire larger fields of view on mineralized bone tissue (Schneider et al., 2011). Therefore, so far, FIB-SEM studies of bone were designed to focus on either the collagen organization or the LCN and could not allow for investigation of both.

A relatively new technique, plasma focused ion beam (PFIB-SEM) presents many advantages for capturing large scale volumes with high resolution. A PFIB is a dual beam FIB-SEM instrument that uses a Xenon (Xe^+^) plasma ion source to achieve faster milling rates than traditional gallium-based FIB-SEM technology. This ultimately results in larger volumes accessible with the same high resolution images afforded by the SEM (Bassim et al., 2014; Burnett et al., 2016). As outlined in Figure 1, this large-volume accessibility enables more of the osteocyte lacuno-canalicular network and a larger proportion of the tissue to be probed in PFIB-SEM compared to traditional FIB-SEM. However, since both utilize an SEM imaging source, the same nanoscale resolution can be achieved. To our knowledge, PFIB-SEM has yet to be employed in the investigation of bone ultrastructure to microscale structure (also referred to as mesoscale structure), despite its success in characterizing other materials with porous features, including concrete (Burnett et al., 2016)and hydroxyapatite coatings on biomedical alloys (Hu et al., 2017).

**Figure 1.**
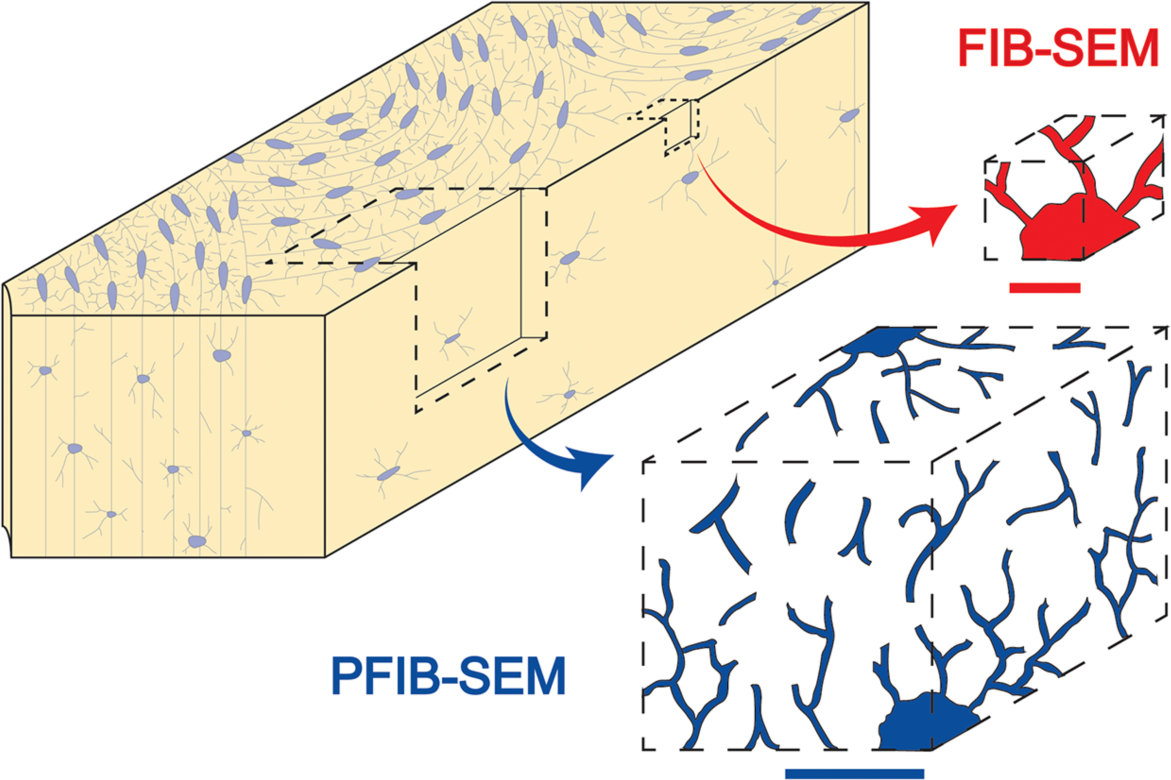
Comparative accessible bone volumes with FIB vs PFIB-SEM with respect to LCN features. Size restrictions in conventional FIB-SEM techniques (red) essentially allow imaging a single osteocyte lacuna in bone tissue due to their inherently small volumes. The PFIB-SEM technique (blue) is able to probe volumes that are an order of magnitude larger than conventional FIB-SEM methods, allowing for high-resolution 3D reconstruction of a larger proportion of lacunae and canaliculi, thus theoretically providing a broader visualization of the LCN network characteristics. Red scale bar: 10µm. Blue scale bar: 50µm.

In this work, we apply PFIB-SEM serial sectioning to investigate demineralized human femoral lamellar bone following state-of-the-art sample preparation protocols. We show, in particular, that the extent of the probed volume allows visualization of a considerably larger portion of the LCN while retaining sufficient spatial resolution to observe the characteristic collagen features. We then show that, without having to go through the burdensome procedure of demineralizing and staining, similar imaging performances can be achieved on mineralized bone. This offers new perspectives which we explore to revisit the mesoscale organization of mineralized microfibrils in human bone highlighting the presence of mineral clusters, a largely overlooked intermediate level of organization between microfibrils and lamellae in human osteonal bone.

## Materials and Methods

### Specimen

The human femur of a 68-year-old male, with no known bone disease, was obtained with institutional ethical approval (HIREB No. 12-085-T) and fixed in a 4% glutaraldehyde (Sigma Aldrich, Missouri, US) solution in a 0.1 M cacodylate buffer for 7 days. The bone was sectioned using a slow speed diamond saw (Buehler Isomet, Illinois, US) under hydrated conditions. A longitudinal section, approximately 2 mm thick, was cut along the length of the femur for demineralization. It was imaged normal to the cutting plane. A transverse section, also 2 mm thick, was extracted for immediate dehydration and embedding, remaining in its naturally mineralized state. This section was imaged normal to the cutting plane. A schematic detailing sample orientation is shown in Figure S1.

### Demineralization and Staining Protocols

In preparation for demineralization, the longitudinally sectioned bone was polished to approximately 200 µm in thickness with 400, 800, 1200, and 2400 grit emery paper and finally a 50 nm diamond suspension on a polishing cloth (Buehler, Illinois, US). The sample was demineralized by immersion in a solution of 5% ethylenediaminetetraacetic acid (EDTA) (Sigma Aldrich, Missouri, US) and 2% paraformaldehyde (PFA; Sigma Aldrich, Missouri, US) in cacodylate buffer, pH 7 until the sample was visibly transparent (approximately 72 hrs). The demineralized bone was rinsed 10 times with deionized water prior to staining. The sample was then pre-stained with Alcian blue using a modified version of a previously published protocol (Reznikov et al., 2014b). 5% Alcian blue in an aqueous solution was heated to 30°C and cooled to room temperature on a hot plate for a total of 10 cycles. The demineralized bone was rinsed 5 times with deionized water prior to subsequent staining. The sample was then stained by successive treatments with osmium tetroxide and thiocarbohydrazide (OTOTO) according to previously described methods (Reznikov et al., 2013).

### Embedding Protocols

Both the demineralized and mineralized tissue were dehydrated in a graded series of ethanol (70%, 80%, 90%, 95%, 100%) for 12 hours each, and further dehydrated in 100% propylene oxide (Sigma Aldrich, Missouri, US). The tissues were gradually infiltrated (25%, 50%, 75%, 100%) with EMbed812 resin (Electron Microscopy Sciences, USA) in propylene oxide (Sigma Aldrich, Missouri, USA). The embedded blocks were then cured in an oven at 60°C. The top surface and adjacent cross-section of the embedded bone were polished with 400, 800, 1200, and 2400 grit emery paper, and a 50 nm diamond suspension on a polishing cloth (Buehler, Illinois, US) to expose the bone in the resin.

### Sample Coating

The polished demineralized and mineralized bone was placed on standard SEM stubs using silver paint. Both the exposed top face and cross-section intended for PFIB-SEM slicing were coated with 5 nm of gold using a Precision Etching Coating System (PECS II) coater (Gatan Inc., CA, USA) to minimize charging effects.

### PFIB-SEM Serial Sectioning: Demineralized Tissue

A Xenon-sourced PFIB microscope (Helios G4 UXe, Thermo Scientific, Hillsboro, USA) equipped with a Schottky field-emission gun SEM was employed. To protect the region of interest, a 63 × 47 × 5 μm^3^ Pt capping layer was deposited on the top surface using a 20 nA and 12 keV ion beam. Brief capping layer experiments (see supplementary methods, Table S1 and S2 and Figure S2) were completed to determine suitable protective layer compositions to minimize curtaining artifacts attributed to ion channeling and preferential milling. Suitable protective layer compositions for demineralized and mineralized tissues were qualitatively determined to be platinum (Pt) and carbon (C), respectively.

Trenches adjacent to the area of interest were milled in order to expose the cross-section and reduce shadowing to detectors and the re-deposition of material. An X-shaped fiducial was milled into the exposed cross-section to allow for post-processing data alignment though translational registration. The electron beam was focused on the exposed cross-section at 2 keV and 1.6 nA under ‘immersion mode’, with a working distance of 5.9 mm, pixel width of 20 nm, a 500 ns dwell time with 2 frame integrations. Imaging was completed with a retractable concentric backscattered detector (CBS). Sequential milling and images were collected using automated Slice and View software (Thermo Scientific, OR, USA), with a 4 nA ion current, 30 keV accelerating voltage, 20 nm slice thickness, and a 4° stage rocking angle, which has been demonstrated to minimize curtaining artifacts (Loeber et al., 2017). A full workflow is visualized in Figure S3. The final cropped volume of the demineralized bone was 45.8 x 40.9 x 8.7 μm^3^.

### PFIB-SEM Serial Sectioning: Mineralized Tissue

Similarly to the demineralized dataset, a protective capping layer 50 μm x 50 μm x 8 μm of C was deposited on the area of interest using ion beam deposition at 60 nA and 12 keV. An X-shaped fiducial was milled into the exposed cross-section to allow for post-processing data alignment. The electron beam was focused on the exposed cross-section at 1.5 keV and 0.8 nA, with a working distance of 2.7 mm, pixel width of 12.5 nm, a 500 ns dwell time, with 8 frame integrations. Imaging was completed with an in-lens detector under backscatter mode. Tomography data was collected using automated Slice and View software (Thermo Scientific, OR, USA), with 1 nA ion current, 30 keV accelerating voltage, 25 nm slice thickness, and a 7° stage rocking angle. The final cropped tomogram of the mineralized bone was 28.6 μm x 25.6 μm x 8.7 μm.

### PFIB Data Reconstruction and Visualization

The tomographic datasets were processed using Dragonfly 4.1 (Objects Research Systems, QC, Canada). Datasets were aligned using a cross-correlation approach and curtain artifacts were removed using the image processing tool box available in Dragonfly. Shadowing effects from the SEM image aligning fiducial were removed on the demineralized images by changing the shading compensation, applying a histogram balance, and also by applying a contrast limited adaptive histogram equalization to the mineralized images. The images were reconstructed, and the osteocyte network was segmented using a U-Net classifier trained with 15 manually segmented images. Segmentations were inspected against the original datasets and minor corrections were implemented manually, including some denoising and selection of missed canaliculi on 2D slices. The machine learning outputs and revised datasets were transformed to a thickness mesh to calculate the canalicular diameter (Figure S4). Cellular organelles and the cell body were segmented manually. Small volumes were extracted from the mineralized dataset to view the mineral morphology, where a Gaussian filter (σ = 0.5) was applied to reduce noise. Orthogonal slices, two-dimensional, and three-dimensional volumes were extracted from these smaller datasets.

It is important to note some geometrical information and the assignment of planes and directions. In our volumes, the XY plane always represents the milling and image plane during acquisition while the YZ and XZ planes are reconstructed after slice registration. By this convention, Z is always the thickness of the serial sectioning dataset. A clarification of the sample geometries from the bulk tissue are clarified in Figure S1.

## Results and Discussion

### Advantages of PFIB-SEM for the visualization of the LCN

One of the main advantages of PFIB-SEM serial sectioning is the large volumes accessible with high resolution which enable exploration of the osteocyte and the LCN. A section of demineralized and stained lamellar bone was reconstructed into a volume measuring 45.8 x 40.9 x 8.7 µm^3^ with an isotropic voxel size of 20 nm (Figure 2A), representing the largest volume of human bone sectioned in a FIB-SEM instrument to this day (see summary of several FIB-SEM experiments in Table S3). Shown alongside the reconstructed volume are orthogonal planes (Figure 2B,C,D), where key features are the osteocyte in the top-left and its surrounding lacuna. While a boundary between the osteocyte and the lacuna is indistinguishable, perhaps due to limited sample preparation, some distinct membrane-bound cellular organelles were present and segmented of which, the nucleus is most prominent (Figure 2E). In all slices, the canaliculi are clearly resolvable, primarily as relatively circular features with a bright boundary, due to osmium tetroxide staining, in the XY plane (arrows Figure 2F) or as oblong, branch-like features in YZ and XZ (Figure 2B, D). While a separate slice in Figure 2G highlights the capability to image the distinct collagen banding patterns also with PFIB-SEM which is a hallmark of most FIB-SEM serial sectioning performed on demineralized bone focused on fibril orientation (Reznikov et al., 2014b, 2013).

**Figure 2.**
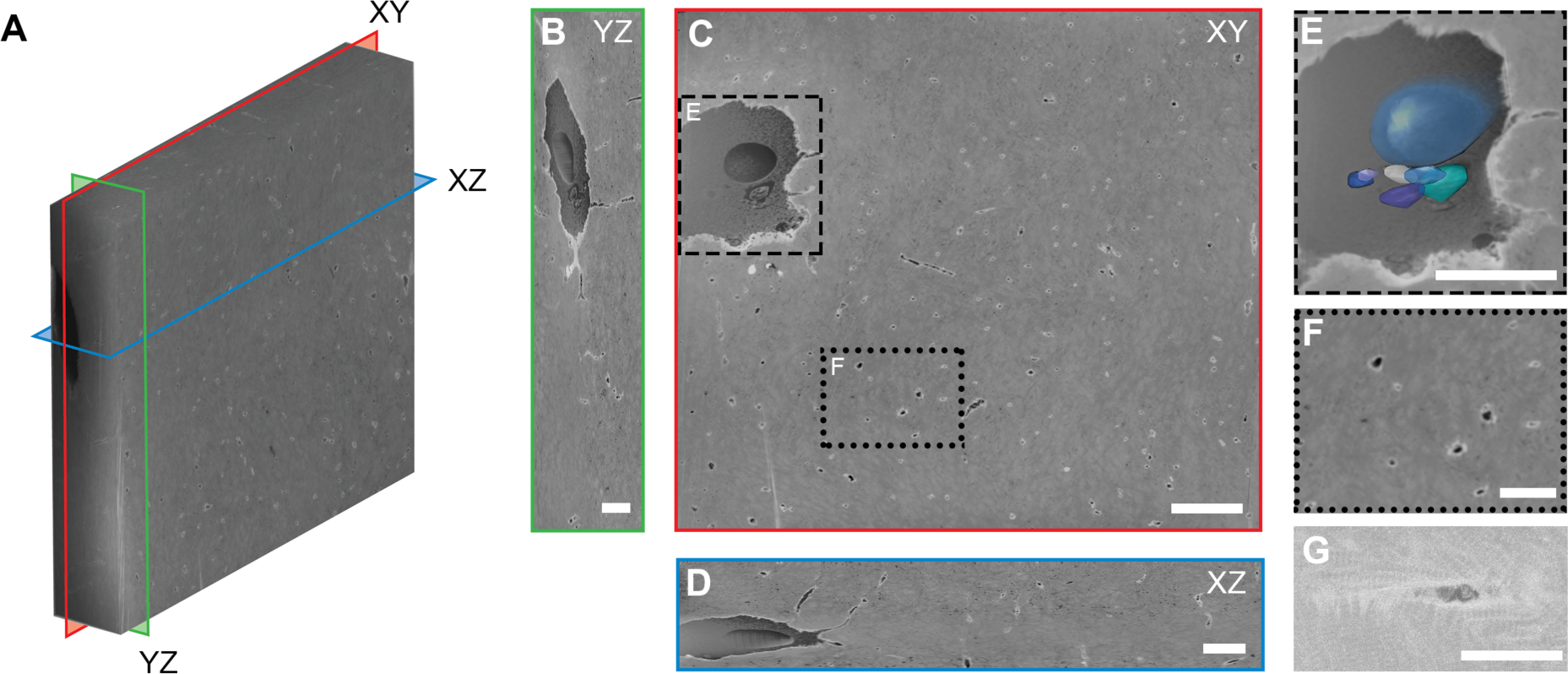
PFIB-SEM of demineralized lamellar bone. (A) 3D reconstruction of demineralized bone with an isotropic voxel size of 20 nm and total volume measuring 45.8 μm x 40.9 μm x 8.7 μm. YZ (green), XY (red), and XZ (blue) orthogonal planes are annotated to show the location of the images in B, C, and D, respectively. (B) An image extracted from the YZ plane of the reconstructed volume, with osteocyte visible. Scale bar: 2 µm. (C) An image of the XY plane. In this view, canaliculi appear as bright circles (inset F). An osteocyte is resolved within the dataset (inset E). Scale bar: 5 µm. (D) An image extracted from the XZ plane of the reconstructed volume. Scale bar: 3 µm. (E) Inset of (C), showing the osteocyte within its lacuna with higher magnification, segmented cellular organelles are overlaid on the image. Scale bar: 5 µm. (F) Inset of (C), where white arrows point to canaliculi in cross-section. Due to staining, their circular periphery appears bright white. Scale bar: 2 µm. (G) A higher magnification image from another demineralized area, showing characteristic collagen banding around a canaliculi. Scale bar: 1 µm.

The segmentation of the LCN from this demineralized dataset is shown in Figure 3A and B, with segmented cellular organelles in C and D. Here, the canaliculi appear to stretch outwards from the osteocyte in all directions, with many connections also observed stretching up from the bottom right-hand side of Figure 3A. Indeed, another osteocyte was located just outside of the volume captured in the tomogram, but visualized during set-up (shown in Figure S5). Therefore, the center of the volume approximately represents the meeting zone of the canaliculi from at least two osteocytes. The canalicular diameter in this segmentation was 346 ± 146 nm (Figure 3E). The volume and segmented LCN of the demineralized sample are available in Supplementary Movie 1.

**Figure 3.**
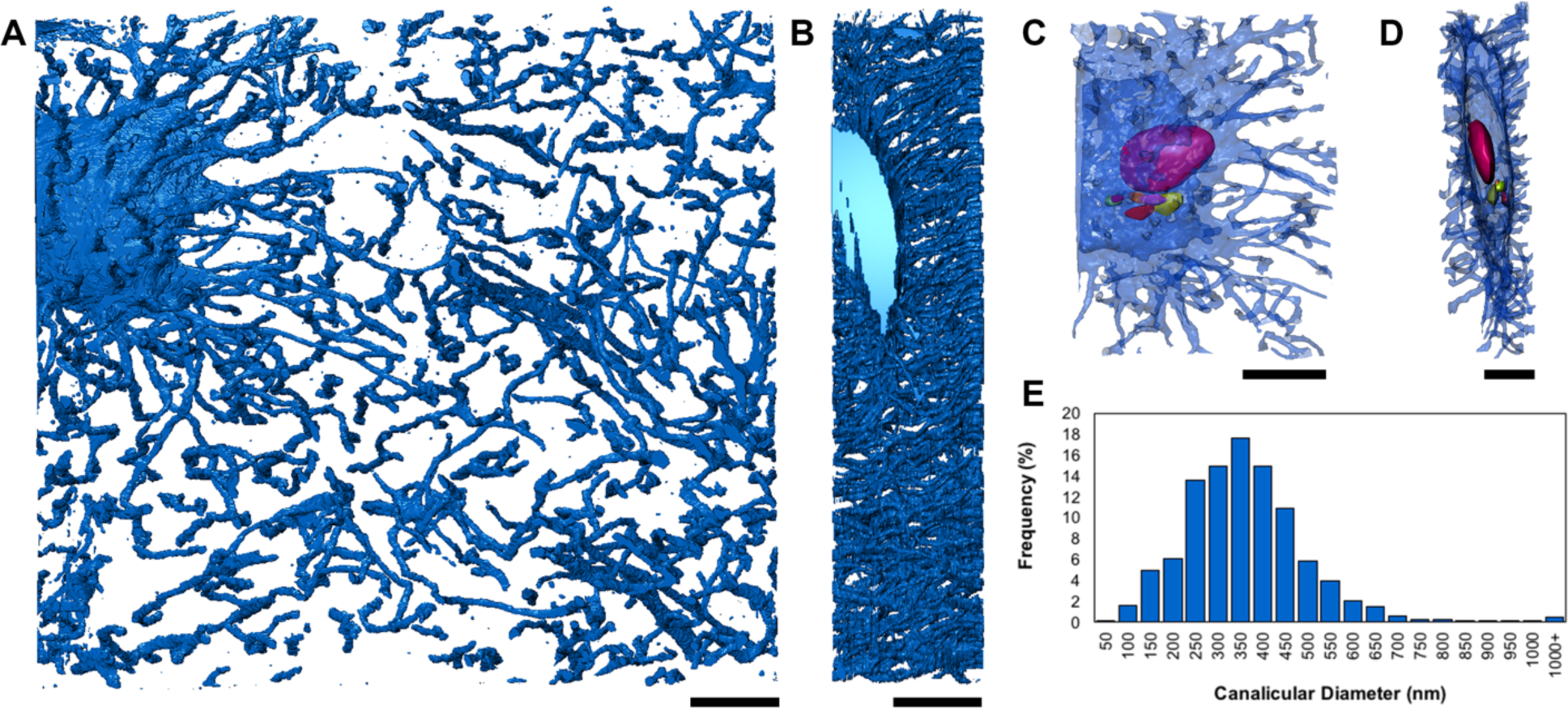
LCN network in demineralized lamellar bone. (A) XY projection of the segmented LCN where canalicular extensions span the volume. More canaliculi appear near the osteocyte cell in the top left corner, and near the bottom right corner of the volume, which is confirmed to be proximal to a second osteocyte that was not captured in the PFIB-SEM dataset (Figure S5). Scale bar: 10 µm. (B) The segmented LCN from the YZ projection, which shows connections spanning the depth of the reconstruction. Scale bar: 5 µm. (C) A segmentation of the osteocyte and nearby extending canaliculi with resolvable cellular organelles segmented (multi-coloured) in 3D. Scale bar: 5 µm. (D) The cell from the YZ plane showing the 3D distribution of cellular organelles. Scale bar: 3 µm. (E) A frequency histogram of canalicular diameter from the segmentation. The average canalicular diameter is 346 ± 146 nm.

Unlike other FIB-SEM of human bone, we also show that mineralized and unstained cortical bone can be analyzed using PFIB-SEM. A slightly smaller 3D volume measuring 28.6 x 25.6 x 8.7 µm^3^ with an anisotropic voxel size of 12.5 x 12.5 x 25 nm is shown in Figure 4A. Similarly, the reconstructed volume is shown alongside its orthogonal planes (Figure 4B,C,D), where the XY plane was acquired during imaging, and the YZ and XZ planes created after image registration. While PFIB can mill extremely large volumes, the precise control of slice thickness well-resolved features in all planes. By avoiding laborious demineralization and staining processes, the resultant volumes and cross-sections show much different contrast. The bright white to light-grey contrast is representative of heavy matter, in this case the mineral, while any empty space, cellular processes, or light elements, like collagen, appear black. An osteocyte with its outer membrane in its lacuna is visible in the bottom left of the section (Figure 4E), although cellular processes are not. These cellular features have been well characterized using traditional FIB-SEM of osteocytes in trabecular bone (Hasegawa et al., 2018, 2017) and near atomic resolution using TEM tomography (Kamioka et al., 2012). Canaliculi are visible as black circles (arrows Figure 4F) in the XY plane and appear as oblong shapes or tubular features (Figure 4B, D) that span the entirety of the YZ and XZ plane.

**Figure 4.**
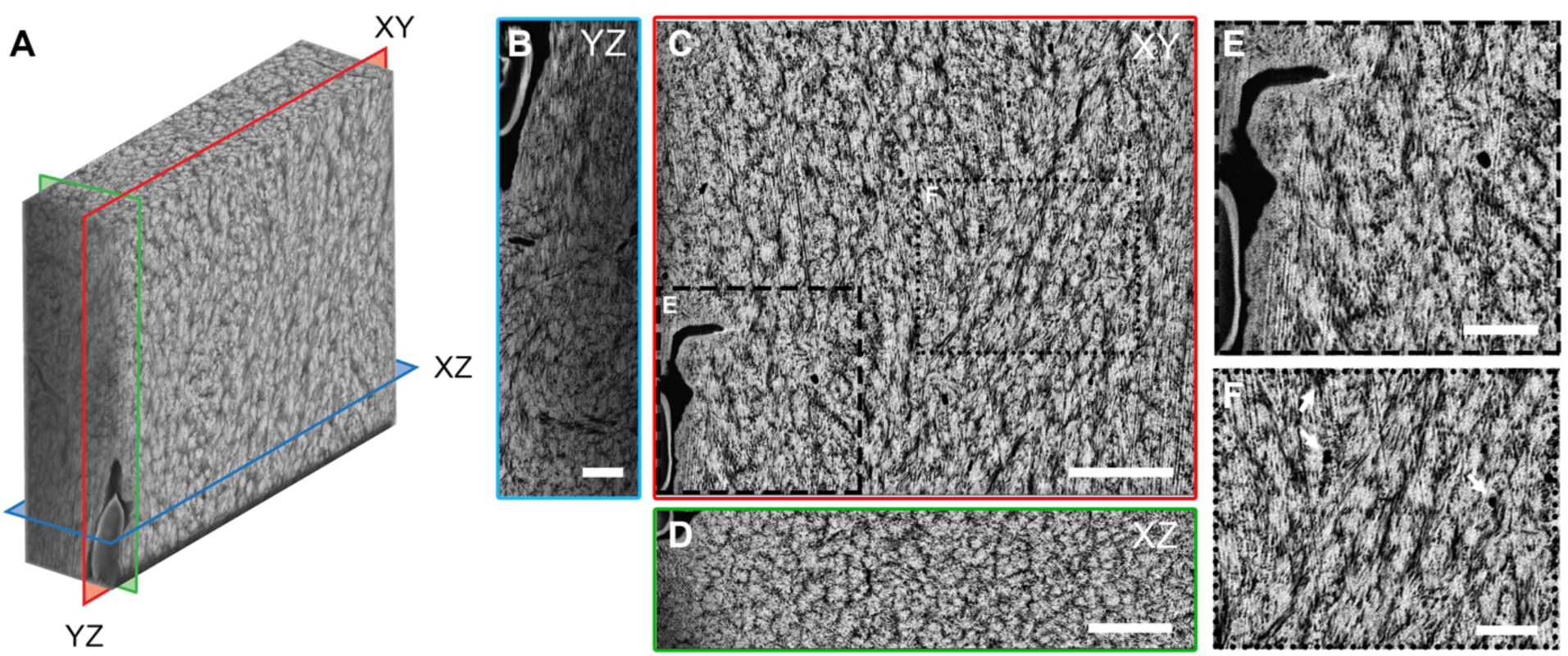
PFIB-SEM of mineralized lamellar bone. (A) 3D reconstruction of mineralized bone with an isotropic voxel size of 25 nm and total volume measuring 28.6 μm x 25.6 μm x 8.7 μm. YZ (green), XY (red), and XZ (blue) orthogonal planes are annotated to show the location of the images in B, C, and D, respectively. Mineral appears light grey/white, while canalicular networks, porosity and collagen all share black contrast. (B) An image extracted from the YZ plane of the reconstructed volume, note an osteocyte in the top left corner and dark canaliculi traversing the view on oblique angles. Scale bar: 2 µm. (C) An image of the XY plane. In this view, the mineral component of bone appears bright, while canaliculi appear as black round circles (inset F). An osteocyte is present in the bottom left (inset E). Scale bar: 5 µm. (D) An image extracted from the XZ plane of the reconstructed volume, corresponding to the transverse anatomical plane. Scale bar: 3 µm. (E) Inset of (C), with osteocyte (bottom left) showing a clear membrane (light grey) and the LCN, which appears black. Cell processes were not visible within canaliculi extending from this osteocyte. Mineral appears finer near the canaliculi periphery, but may be due to shadowing. Scale bar: 2 µm. (F) Inset of (C), where white arrows point to circular canaliculi. Note that due to the absence of staining, the canaliculi in this case appear black. Scale bar: 2 µm.

The segmentation of the LCN network from this mineralized dataset is shown in Figure 5A and B, with the cellular membrane segmented in C and D. Most of the canaliculi appear co-aligned in the Z-direction (Figure 5B) within this volume, with minimal branching across the volume. It is well known the canalicular density can vary strongly within different regions of a bone sample. Also, in osteons, osteocytes are flattened in the tangential plane (in the lamellar plane) with the highest density of canaliculi oriented radially. Recall, that the samples were cut in orthogonal directions and planes were assigned such that the XY plane was the milling and image plane in PFIB-SEM (Figure S1). Based on the images of segmented canaliculi, it is most likely that the demineralized XY plane falls radially with respect to an osteon, while the mineralized XY plane likely falls tangentially with respect to an osteon. The average canalicular diameter in the mineralized bone was 284 ± 82 nm (Figure 5E), still in line with values reported by other techniques but a narrower range in comparison to the diameters reported for the demineralized bone above (346 ± 146 nm). This difference could either be explained by a loss of mineral at the canalicular surface during demineralization, or to the difficulty to precisely define this interface due to a relative diffuse staining at the canaliculi surface of demineralized bone. Interestingly, 33 % of the values fall in a relatively narrow range of 300-400 nm for the demineralized sample while 58 % of the canaliculi diameters are slightly more broadly distributed within 250-450 nm. For the mineralized sample, 50% are between 300-400 nm and 73% between 250-450 nm. It is important to note, however, that a precise quantification of canaliculi diameter strongly depends on the segmentation procedure. Figure S4 highlights differences in segmentation, when meticulously corrected manually (shown in Figure 3 and 5), or when segmentation is completely relaxed and automated. The volume and segmented LCN of the mineralized sample are available in Supplementary Movie 2.

**Figure 5.**
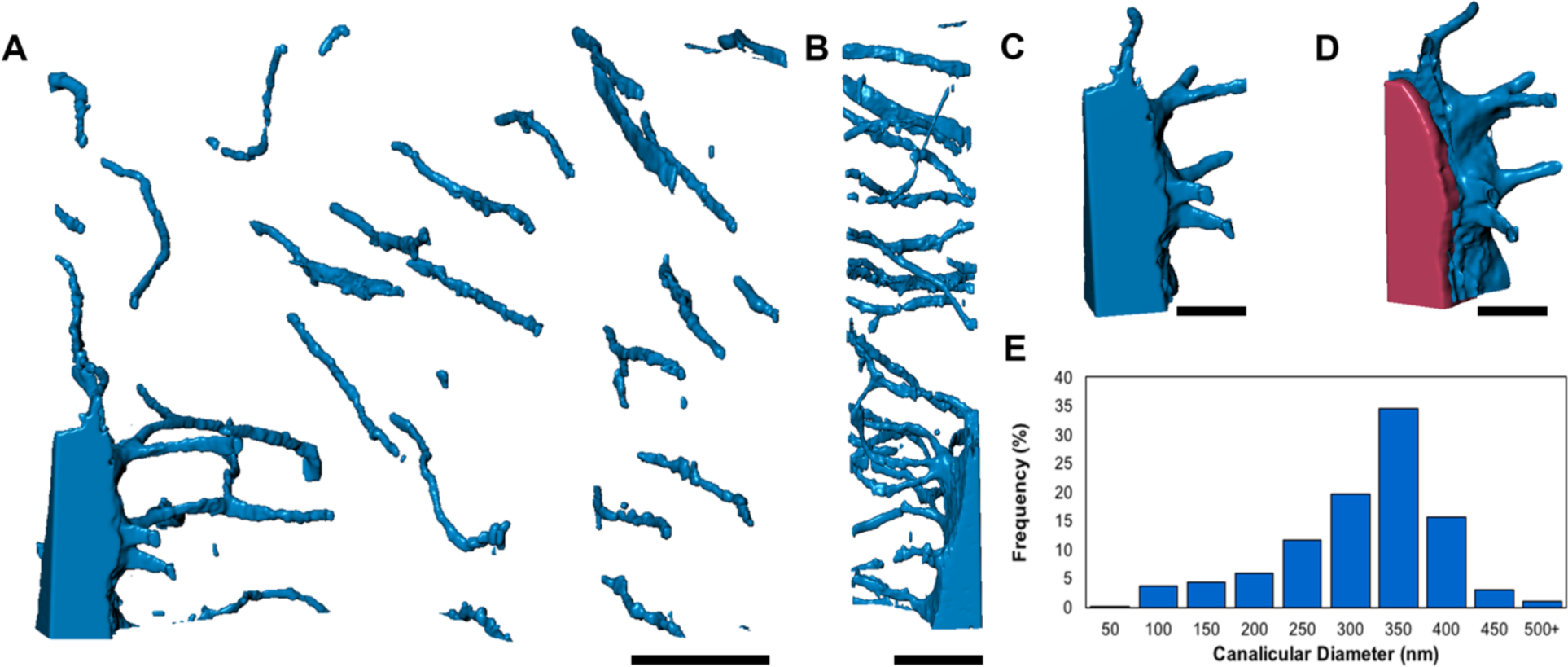
LCN network in mineralized lamellar bone. (A) XY projection of the segmented LCN where long canalicular connections appear to extend primarily along the Z-direction of the reconstruction. This corresponds to along the XZ or transverse plane, therefore likely part of a network that would span osteonal layers. Part of an osteocyte is present in the bottom left corner of the segmentation. Scale bar: 5 µm. (B) The segmented LCN from the YZ view, which shows co-aligned connections spanning the depth of the reconstruction, with most being aligned in this direction. Scale bar: 5 µm. (C) A segmentation of the lacunae and extending space that (D) the osteocyte cell (pink) resides in. Scale bar 2 µm. (E) A frequency histogram of canalicular diameter from the segmentation. The average canalicular diameter is 285 ± 82 nm.

Osteocytes or LCN have been studied with a variety of techniques. Functional studies mostly rely on optical microscopy, due to the large variety of available fluorescent dyes that allow targeting specific cellular components and biomarkers (Ciani et al., 2009; Sano et al., 2015). Confocal and non-linear two-and three-photon microscopy have also been used to study the LCN characteristics and have the additional advantage of allowing large fields of views (Genthial et al., 2017; Kerschnitzki et al., 2011; Repp et al., 2017). However, all these methods suffer the classical diffraction limitation and the spatial resolution is typically limited to ∼ 200 nm laterally and ∼ 500 nm axially, such that the smallest canaliculi are out of reach, thus complicating network connectivity analysis. Higher resolution afforded by SEM was used to report canaliculi diameters in a range of 259 ± 129 nm on 2D sections of mice humeri (You et al., 2004). Similarly, atomic force microscopy (AFM) studies report canaliculi diameters of 426 ± 118 nm on 2D sections of cortical bovine tibia (Lin and Xu, 2010).

To overcome the limits of dimensional quantification on 2D observations, pseudo-3D visualization is often performed with SEM using resin-embedded and acid-etched bone replica (Bonewald, 2011; Shah and Palmquist, 2017). In 3D, two partial osteocyte lacunae and their junctions in murine bone were probed by traditional FIB-SEM (Schneider et al., 2011). However, the roughly 20 µm wide view did not capture a full lacuna, nor an osteocyte maintained within. While osteocytes and individual organelles were not classified in this work, as was not the aim, the feasibility to probe cellular organelle structures opens a potential avenue for further PFIB-SEM applications.

X-ray tomography is also frequently used to assess various characteristics of the LCN. However, reaching sufficient resolutions to capture the smallest canaliculi still remains a challenge. Canaliculi with diameters of 320-390 ± 120 nm were reported for human cortical femurs analyzed by X-ray phase contrast synchrotron nanotomography (SR-nanoCT) (Varga et al., 2015), but such measurements currently provide typical resolutions of 100-200 nm in the best cases (Varga et al., 2016). These works characterize the osteocyte network and LCN of mineralized bone tissue, and reveal some texture that is attributed to the collagen banding pattern (Peyrin et al., 2014). However, while a larger volume is obtained, even using a synchrotron source, voxel sizes ranged from 50 to 130 nm (Wittig et al., 2019) to 280 nm (Peyrin et al., 2012), double to a full order of magnitude larger than the 25 nm obtained herein. Conversely, the advancements in image processing for X-ray data are significantly more developed than the nascent field of PFIB, and reconstructed and denoised to reveal fine details of bone structure (Dong et al., 2014; Pacureanu et al., 2013, 2012). Future PFIB-SEM evaluation of bone structure and LCN should focus on further optimization of acquisition parameters as well image processing techniques to reveal the structural and biological linkage in bone. Ultimately, PFIB-SEM could advantageously provide accurate quantification of local regions of the LCN if coupled with other less resolved methods which provide a visualization of larger portions of the network.

### Evidence of ubiquitous microscopic mineral clusters in bone tissue

In addition to the features associated with the presence of osteocytes, the PFIB-SEM images of mineralized bone in Figure 4 exhibit much more contrast fluctuations than can be found in the demineralized volume of Figure 2. This contrast must logically be associated to mineralization fluctuations in the tissue. Upon closer inspection of the mineralized data, this contrast takes the form of regular bright patches surrounded by a darker border scattered throughout the imaged volume with dimensions in the micron range. Moreover, when viewed in two orthogonal directions, those patches appear smaller and more circular in the transverse direction (Figure 4D) and more elongated longitudinally (Figure 4B). To analyze those patches, five sub-volumes (cubes with side length of 3 µm) were selected along a random diagonal direction (Figure 6 and Figure S6). These highlight more clearly the shape and orientation of those mineral clusters: they appear more circular in the XZ plane (transverse to osteon), irrespective of the ROI location in the sample and are more elongated in the orthogonal planes, especially YZ where the smallest features have dimensions compatible with mineralized collagen microfibrils aligned in the longitudinal plane. Extracting one representative mineral cluster and showing all its orthogonal planes and as a 3D reconstruction (Figure 7), the shape of these structures in 3D more closely resembles a flattened prolate ellipsoid, marquise, or 3D elongated diamond, than a perfect sphere, the elongation direction being the osteonal axis. The mineral clusters measure approximately 1 µm in width and 2-3 µm in length. These are shown across the whole XZ plane in Supplementary Movie 3 and in more detail in Supplementary Movie 4.

**Figure 6.**
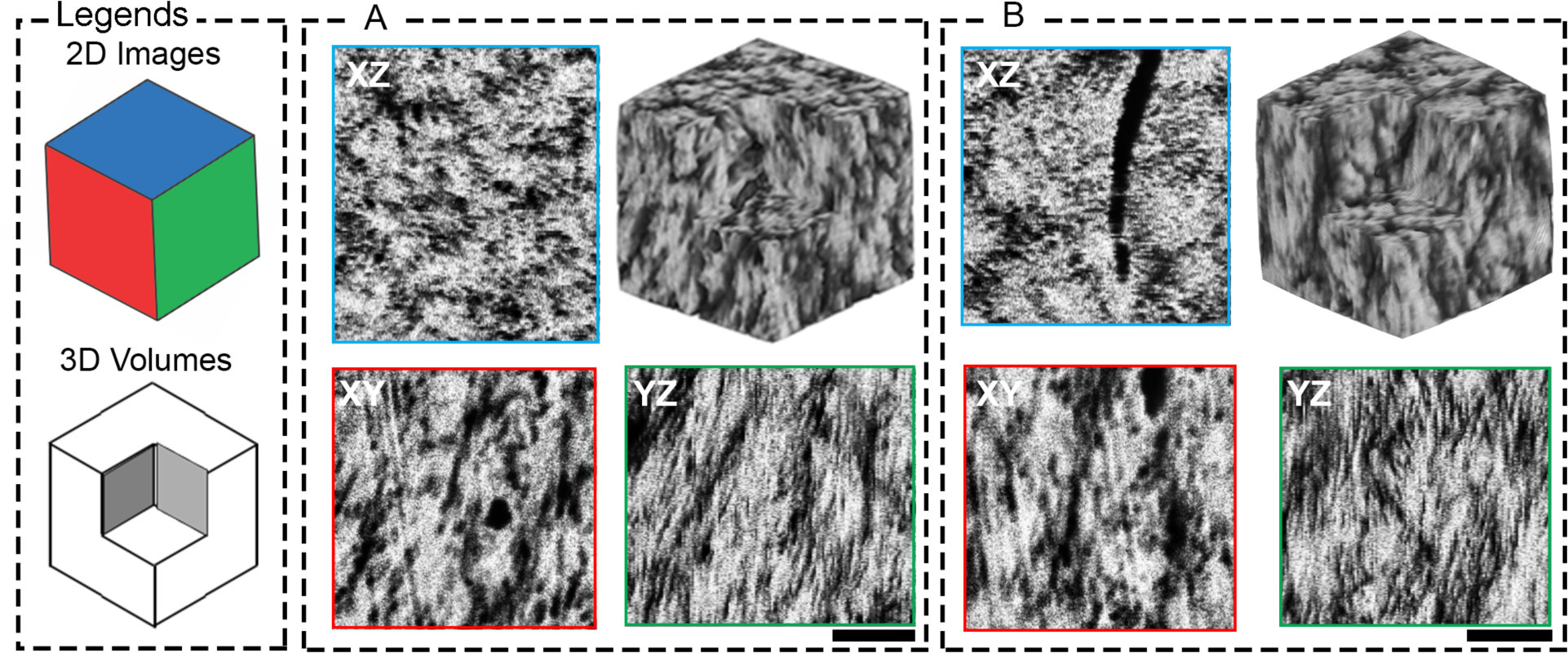
Exploring mineral and collagen orientation in 3D. The PFIB volume is cropped into smaller volumes (A,B) measuring 3 µm each side to explore the orientation of mineral and collagen fibrils. The transverse planes (XZ) show clear rosette shapes, while the YZ plane along the long axis of the femur, shows collagen fibrils in plane, denoted by their characteristic banding pattern. XY planes show the faint outline of diamond, or marquise shapes.

**Figure 7.**
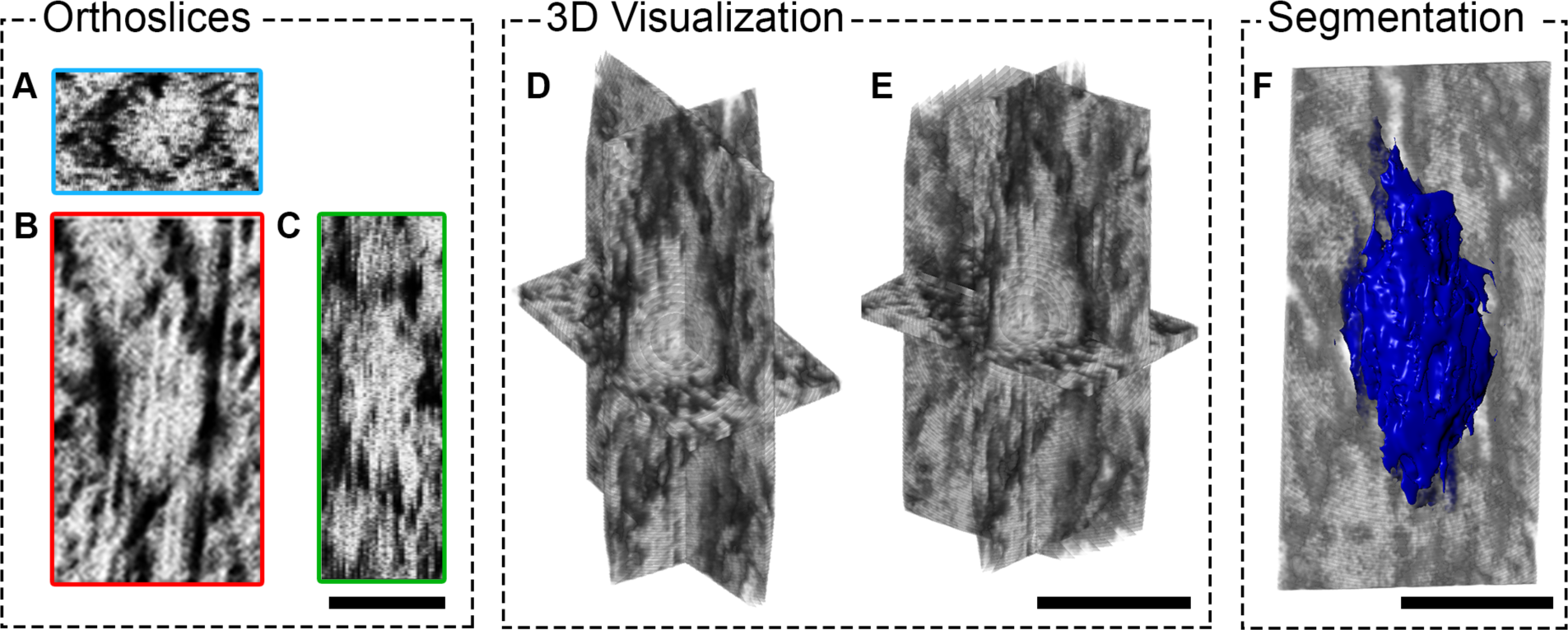
Mineral cluster morphology in 3D. A representative mineral cluster from PFIB-SEM of the mineralized sample has been isolated to probe its shape both by separated (A,B,C) and intersecting orthogonal planes (D,E). In (A) the transverse slice, perpendicular to the long axis of the femur, is given by plane XZ. Here, a clear rosette shape is visible also discernible from the top-down tilted view in (E). Segmentation of the 3D shape in (F) confirms its likely shape as a prolate ellipsoid. All scale bars: 1 µm

Similar mineral clusters have been identified on the floor of osteocyte lacunae (Shah et al., 2016) and more recently, at the apex of rat calvarial sutures (Shah et al., 2020) by exposing bone to deproteinization protocols. In Shah *et al*. the shape has been identified as a marquise, which is not uniform throughout bone, but rather changes to a less organized marquise moving away from the bone mineralization front, suggesting that this shape is a transient early form of mineral deposit (Shah et al., 2020). The same general shapes, identified as prolate ellipsoids, have been shown in the turkey tendon (Zou et al., 2019), and over small sections of human bone, e.g. Movie S1 in Reznikov et al. (Reznikov et al., 2018).

We note that this structure is ubiquitous throughout an entire 45 µm section of lamellar bone and represents a hierarchical level of mineral organization above the single collagen fibril level of 100 nm in diameter and below the typical dimensions of an osteon lamella of 5-7 µm. It also shows that this structure is not a member of the “disorganized” sub-lamella motif (Reznikov et al., 2013) associated with canaliculi. The two earlier papers citing ellipsoidal mineral shapes have also postulated that the presence of a cross-collagen fibril mineralization motif may be possible (Reznikov et al., 2018; Zou et al., 2019). Our findings on the microscale, depicting mineral clusters that span roughly 1 µm in width and 2-3 µm in length across over 45 µm in width clearly indicate that mineral is indeed associated with more than one collagen fibril, confirming earlier hypotheses that tissue mineralization occurs with high spatial regularity over large distances past the single collagen fibril level.

Seminal works focusing on early bone formation by Bonucci *et al*. and Bernard and Pease outline similar spherical shapes as “calcification loci” or “round bodies” between collagen fibrils in osteoid (Bernard and Pease, 1969; Bonucci, 1971). These are thought to elongate, firstly unrelated to collagen banding, then lastly become associated with the fibril interior, or to coarsen from calcification loci to bone nodules which coalesce to form all mineralized tissue (Bernard and Pease, 1969; Bonucci, 1971). These findings are supported by cryo electron microscopy studies, which observe the accumulation of mineral on the exterior of the fibril in *in vitro* conditions (Nudelman et al., 2013). Similar spherical particles ‘calcospherulites’ have been postulated to be related to the mineralization front only (Midura et al., 2008) and early bone mineralization from *in vitro* culture of MC3T3-E1 osteoblast cells show comparable spherical particles (mineralization foci), and also notes these in mouse calvarial bone with osteopontin closely associated with their margin (Addison et al., 2014).

### The mineral: linking to the nanoscale

Viewing the mineral clusters in the transverse (XZ) plane, means that we view a section perpendicular to the long axis of the osteon, closely aligned along the femur axis in our case and, presumably, of the collagen fibrils. In such an orientation the circular shape is fully consistent with previously observed features in high-resolution TEM (Figure 8) in human bone, termed “rosettes” (Grandfield et al., 2018), “lacy” pattern (Reznikov et al., 2018) or otherwise unnamed in the alveolar bone of minipigs (Maria et al., 2019). Dispersed between some rosettes are clear black pores marking the canaliculi, and a substantial portion of black empty space surrounds each rosette. Upon closer inspection of Figure 8B, a unique feature of each rosette is its mineral-dense center (Supplemental Movie 5).

**Figure 8.**
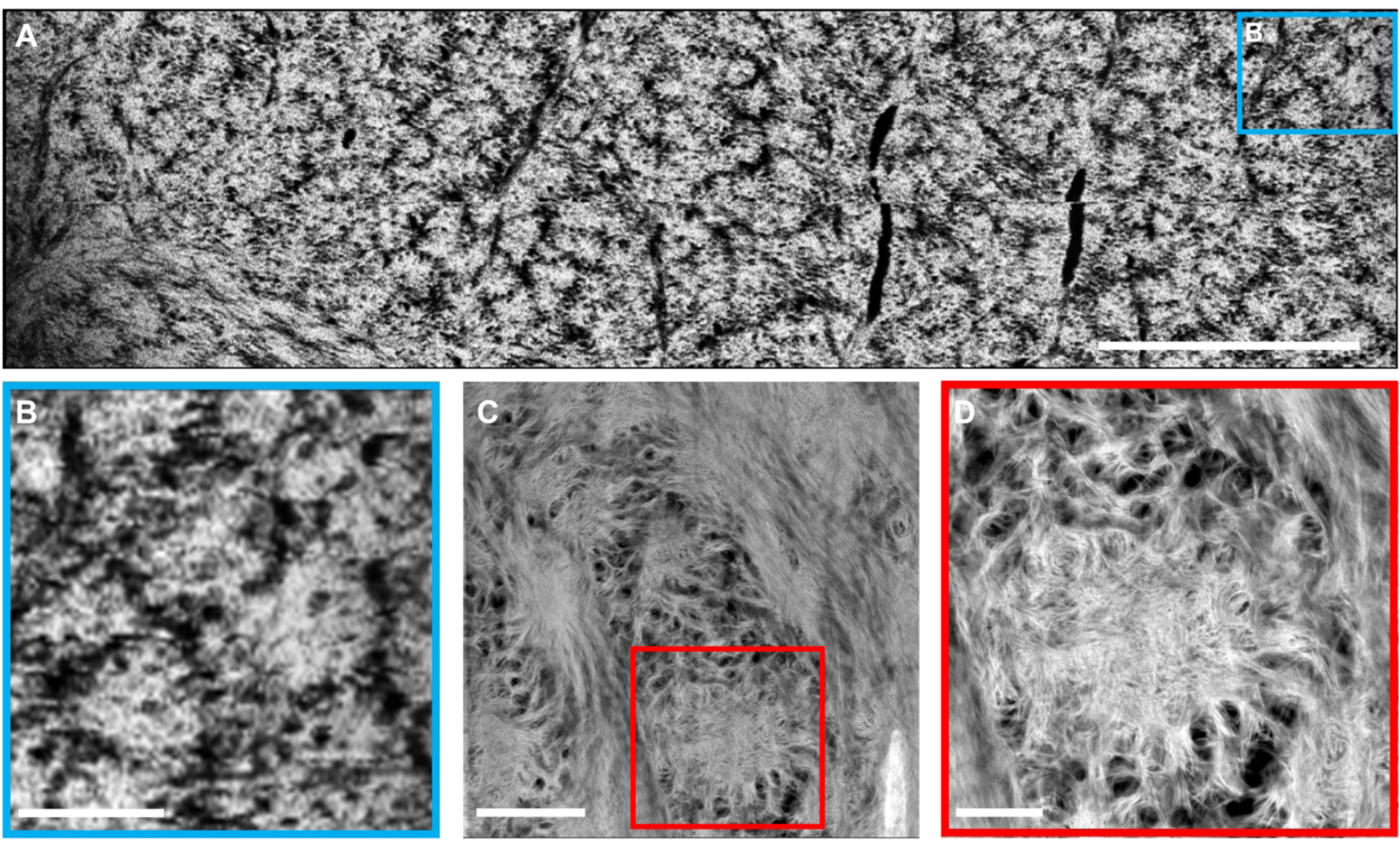
Mineral clusters linked to the nanoscale. A transverse slice (perpendicular to the long axis of the femur) from the mineralized PFIB-SEM volume is shown in (A) with higher resolution inset in (B). Scale bars: 5 µm and 1 µm, respectively. The mineral appears in rosette shapes across the entirety of the section. Black space surrounds the clusters, or takes the form of black lines represent the canalicular network running through the section. A lacuna is located near the bottom left corner, but not shown on this slice. A side-by-side comparison with STEM-HAADF images of human femoral bone (C) with higher magnification inset (D), reprinted with permission from (Grandfield et al., 2018), shows that the black space in PFIB surrounding the rosettes is comprised of mineral plates, wrapping around circular voids, representing collagen, while the central portion remains densely mineralized. Scale bars: 500 nm and 200 nm, respectively.

The rosette features noted by (Grandfield et al., 2018) in large TEM sections, are reproduced in higher resolution images in Figure 8C and D for comparison. With the higher resolution and compositional contrast afforded by high angle annular dark field scanning transmission electron microscopy (HADDF STEM), we begin to make sense of the black regions surrounding the rosettes as viewed in PFIB-SEM. The features are outside of the resolution limit of the PFIB-SEM, but with the STEM images, we can see individual mineral plates exist in this space, and are clearly resolved, wrapping around perfectly circular black voids. We postulate that these black voids are one of two features; sub-cellular porosity or collagen viewed in cross-section, and most likely a combination of both.

While many comprehensive reviews cover the hierarchical levels of bone mineralization (Reznikov et al., 2014c; Weiner and Traub, 1992), we should draw attention to some of the competing views on collagen-mineral arrangement, chief of which is the concept of intra versus interfibrillar mineralization, referring to mineral either occupying (mainly the gap zone of) the collagen fibril, or external to the fibril/between fibrils. This conflicting view essentially stems from convincing results reported over decades supporting both observations. For example, X-ray or neutron diffraction peaks can readily be observed at small angles with the same 67 nm periodicity as this exhibited by non-mineralized collagen fibrils of bone or tendon, pointing to intrafibrillar mineralization in the collagen microfibril gap zone (Bigi et al., 1988), as confirmed by TEM (Weiner and Traub, 1989). More recent STEM results obtained with FIB-SEM ultrathin samples suggest, on the contrary, that mineral nanoparticles are quasi-exclusively extrafibrillar (McNally et al., 2012; Schwarcz, 2015). However, Landis *et al*. clearly reported electron microscopy evidence for the presence of both intra and extrafibrillar mineral in turkey tendon more about three decades ago (Landis et al., 1991) which is likely confirmed by our STEM observations of rosettes.

Nevertheless, how this occurs or what this looks like remains elusive. Our images over large volumes with 12.5nm resolution in X and Y and 25 nm resolution in Z, complemented by those from Grandfield *et al*. (Grandfield et al., 2018), suggest the potential for a combination of intra and interfibrillar mineralization. At least a part of the black voids on the periphery, likely represent collagen in cross-section and therefore, interfibrillar mineralization. In fact, since the black voids make up such a large fraction of the sample it is highly unlikely they are all vacant of any material. Collagen, as a light element, will not scatter electrons well in the PFIB-SEM or TEM, meaning that it likely will always appear black in cross-section. The presence of collagen in these cross-sectional voids has been confirmed in TEM sections using elemental analysis (Lee et al., 2019). On the other hand, the central area of the rosettes, viewed in our PFIB and the complementary STEM (Figure 8 B,C,D), suggests that the central portion may be much more densely infiltrated with mineral. Yet, even in these densely mineralized regions, there are still signs of discrete particles albeit less well defined than at the rosette periphery. While PFIB-SEM is still in its infancy, optimization of the voxel size and slice thickness will enable probing this hypothesis in the future. Indeed, with the addition of elemental information by energy dispersive X-ray spectroscopy (EDX) mineral transport in other species has been shown on single slices of a block-face (Kerschnitzki et al., 2016b). Multi-spectral tomography or serial sectioning, i.e. collection of both images and EDX spectra slice-by-slice, could confirm both nanoscale structures and mineral locations in the future.

### Study Limitations

While this paper provides an exciting proof-of-concept for using PFIB-SEM serial sectioning to analyze bone, we are aware that there are some limitations to our study design. Firstly, we recognize that the demineralized and mineralized datasets represent a single data point and from different orientations in bone – along and transverse to the long axis of the femur. Nevertheless, the findings presented in a single dataset are representative of volumes larger than several datasets of traditional FIB-SEM. Of course, our future work will probe a wider range of samples, both anatomical location and species. We do note that specimens were prepared by dehydration and embedding, rather than cryogenic FIB-SEM. Future possibilities for cryo-PFIB-SEM exist. Nevertheless, our findings are in line with structures shown in cryogenic-FIB-SEM, and embedded FIB-SEM published by others (Table S3). The volumes probed in this study, while larger by an order of magnitude than any other FIB-SEM dataset on bone, are still on the small scale for what is possible with PFIB-SEM. Larger volumes, capturing several osteocytes and their connections, will indeed be interesting to probe in the future, bringing PFIB-SEM to the level of investigations carried out with X-ray techniques. Lastly, in our work, we’ve analyzed a skeletally mature bone specimen, though it is true that the age of the osteon and specific lamella probed is unknown. While we anticipate the prolate ellipsoid or marquise structure is not associated only with mineralization fronts, future work comparing primary and secondary osteons would be interesting to shed light on whether this is a permanent or transient structure in lamellar bone.

## Conclusions

PFIB-SEM is a promising technique to probe both the nano and microscale hierarchy of bone. Herein, we demonstrate that PFIB-SEM serial sectioning can achieve volumes between 2-4 times larger than traditional gallium-based FIB-SEM, while maintaining state-of-the-art resolution of collagen microfibrils. The large volume capabilities of this technique enabled an approach to begin probing the LCN and microfibrils simultaneously. Moreover, this work demonstrates that analyzing mineralized bone, as opposed to demineralized bone, is essential for making claims on collagen-mineral arrangement and bone hierarchical structure in general. As suggested by others in human bone (Reznikov et al., 2018), rat bone (Shah et al., 2020) and turkey tendon (Zou et al., 2019), we confirm that the mineral in cortical bone takes on a ellipsoidal, marquise, or prolate ellipsoidal shape. Our large scale imaging confirms that these clusters are indeed densely packed across lamellar bone and not isolated events. The central cross-section of these ellipsoids represents the “rosette” features described previously (Grandfield et al., 2018). It is therefore clear that the organization of mineral is not solely restricted to its inter-versus intrafibrillar arrangement with collagen, but depends on coordination across several collagen microfibrils. We suggest that the presence of inter-fibrillar mineral is likely on the exterior of these clusters, while intrafibrillar mineral appears to be a possibility in the interior. Further work is needed to probe this hypothesis and investigate how this arrangement changes across osteons, and of course, in other types of bone.

## Acknowledgements

We gratefully acknowledge Dr. Xiaoyue Wang for assistance with bone demineralization, staining and preliminary imaging. Sample preparation was conducted at the Faculty of Health Sciences Electron Microscopy Facility at McMaster University, with assistance by Marcia Reid. Electron microscopy was performed at the Canadian Centre for Electron Microscopy, a facility supported by NSERC and other governmental agencies. We extend our gratitude to Dr. Natalie Reznikov at Object Research Systems (ORS, Montreal) for Dragonfly software support for data visualization and processing of the demineralized dataset.

## Funding Sources

This work was supported by the Natural Sciences and Engineering Research Council of Canada (NSERC) (RGPIN-2014-06053), the France-Canada Research Fund (FCRF-2018-Grandfield), and the Ontario Ministry of Research, Science and Innovation (Early Researcher Award ER17-13-081). DMB and JD are supported by an NSERC CGS-D and NSERC PGS-D scholarship, respectively.

## Author Contributions

Conceptualization; DMB, AG, KG

Methodology: HY, DMB

Data curation; DMB, JD, HY

Formal analysis; DMB, JD

Funding acquisition and supervision; AG, KG

Writing: original draft; DMB, AG, KG

Writing: review & editing; DMB, JD, HY, AG, KG

## SUPPLEMENTARY INFORMATION

**Figure S1.**
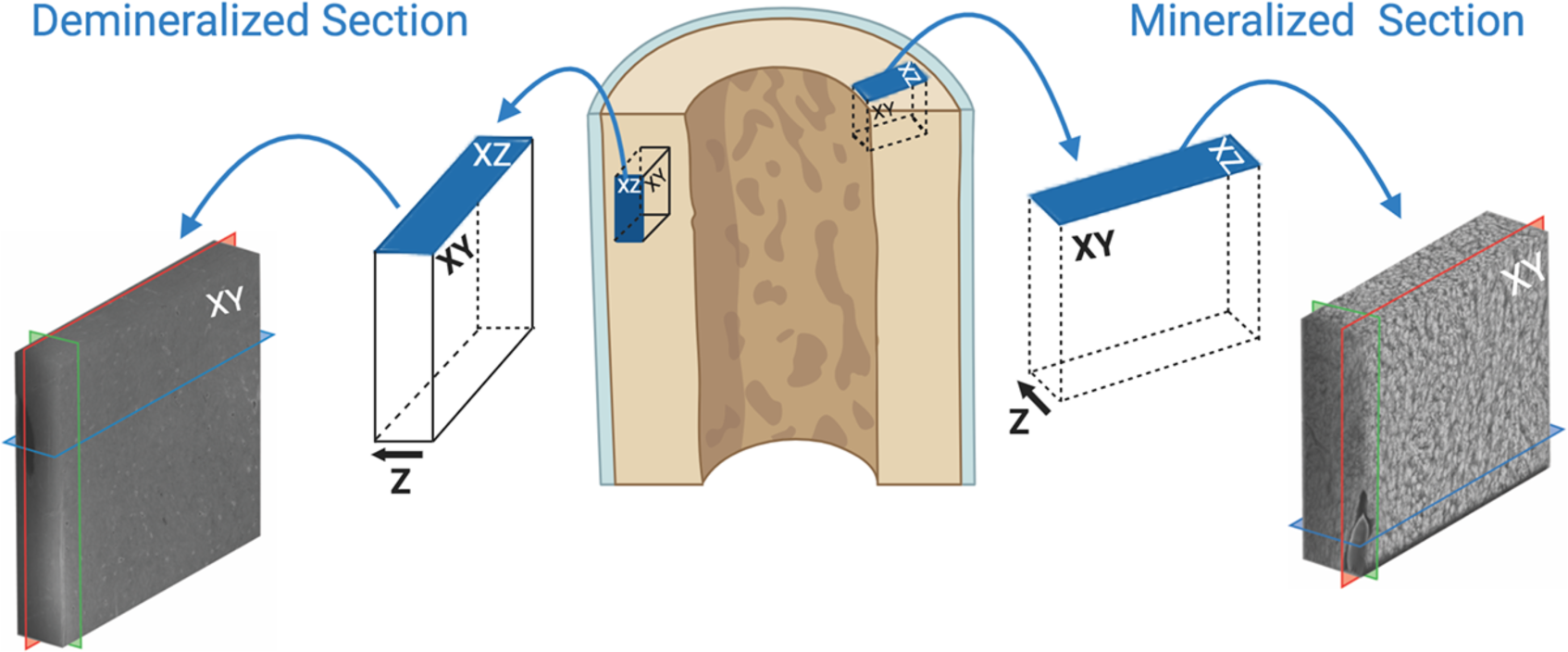
Schematic of sample orientations. The samples were originally cut in the longitudinal (demineralized) and transverse (mineralized) direction. Therefore, the blue planes highlighted above sit on cut surfaces, and represent the top surface viewed in the SEM. In PFIB-SEM, the XY plane is generally assigned normal to the top surface – it represents the surface that is sequentially milled and imaged. The Z direction in PFIB-SEM thereby dictates the direction of sequential milling, and eventually the thickness of the volume. Although the samples were oriented orthogonal to one another, we can see from above that their XY planes both contain the long axis of the bone. Corresponding final tomograms of both from Figures 2 and 4 on the far left and right. Segmented canaliculi in Figure 3 and 5 suggest that the demineralized XY plane was oriented radially with respect to an osteon, while the mineralized XY plane is likely tangential.

### Methods for Capping Layer Composition Experiments

To determine a suitable protective layer against ion beam damage for demineralized and mineralized bone tissue, due to their varying hardness, a variety of protective layer compositions were deposited on the specimens. We explored several options, non-systematically. In the gas injection system, the pure Pt and W deposits are approximately 20% C due to the gas mixture in the gas injection system (Utke et al., 2008). For our purposes, we refer to this as ‘pure’ in Table S1 to avoid confusion. Along with the pure compositions, various flux percentages (%), given as a fraction of the maximum gas output for that element, were explored. After deposition, slices of material were removed using the beam conditions given in Table S2 and the quality of cross-section was assessed qualitatively. A “good” capping layer was determined to have minimal curtaining artifacts by qualitative assessment (Table S1). Our findings (Figure S2 and Table S1) indicate that demineralized bone sliced with a pure Pt capping layer produced the fewest artifacts, while mineralized bone produced the best quality when protected with pure C. These results make sense considering the relative hardness of the capping layers and underlying material, matching a softer Pt coating to demineralized bone, and a harder C coating to mineralized bone. Of course, a larger, more systematic study would be interesting, as most FIB-SEM tomography is conducted with a standard Pt coating.

**Table S1:** Various capping layers were non-systematically investigated to determine optimum composition for minimizing curtaining on demineralized and mineralized bone samples during PFIB-SEM tomography acquisition. The quality of the cross-section, given by a reduction in curtaining artifacts, is ranked qualitatively.

| Quality of Cross Section | Capping Layer Composition |  |  |  |  |
| --- | --- | --- | --- | --- | --- |
|  | Pure C | Pure W | Pure Pt | 80% C to 100% Pt | 90% C to 100% Pt |
| Demineralized Bone | Fair | Good | Good | N/A | N/A |
| Mineralized Bone | Good | N/A | Poor | Fair | Fair |

**Table S2:** Electron and ion beam parameters for cross-sectional milling of demineralized and mineralized bone during protective layer composition experiments. An in-lens detector in backscattered mode was used.

|  | Demineralized Bone | Mineralized Bone |
| --- | --- | --- |
| Protective layer size | 10 x 20 x 5 $\mu\text{m}$ | 20 x 40 x 5 $\mu\text{m}$ |
| Electron beam | 1 keV, 1.6 nA | 1 keV, 3.2 nA |
| Working distance | 4.1 mm | 4.1 mm |
| Ion beam | 1 $\mu\text{A}$, 30 keV | 0.5 $\mu\text{A}$, 30 keV |
| Slice thickness | 15 nm | 100 nm |
| Number of slices | 30 | 5 |
| Rocking angle | 4° | 0° |

**Figure S2.**
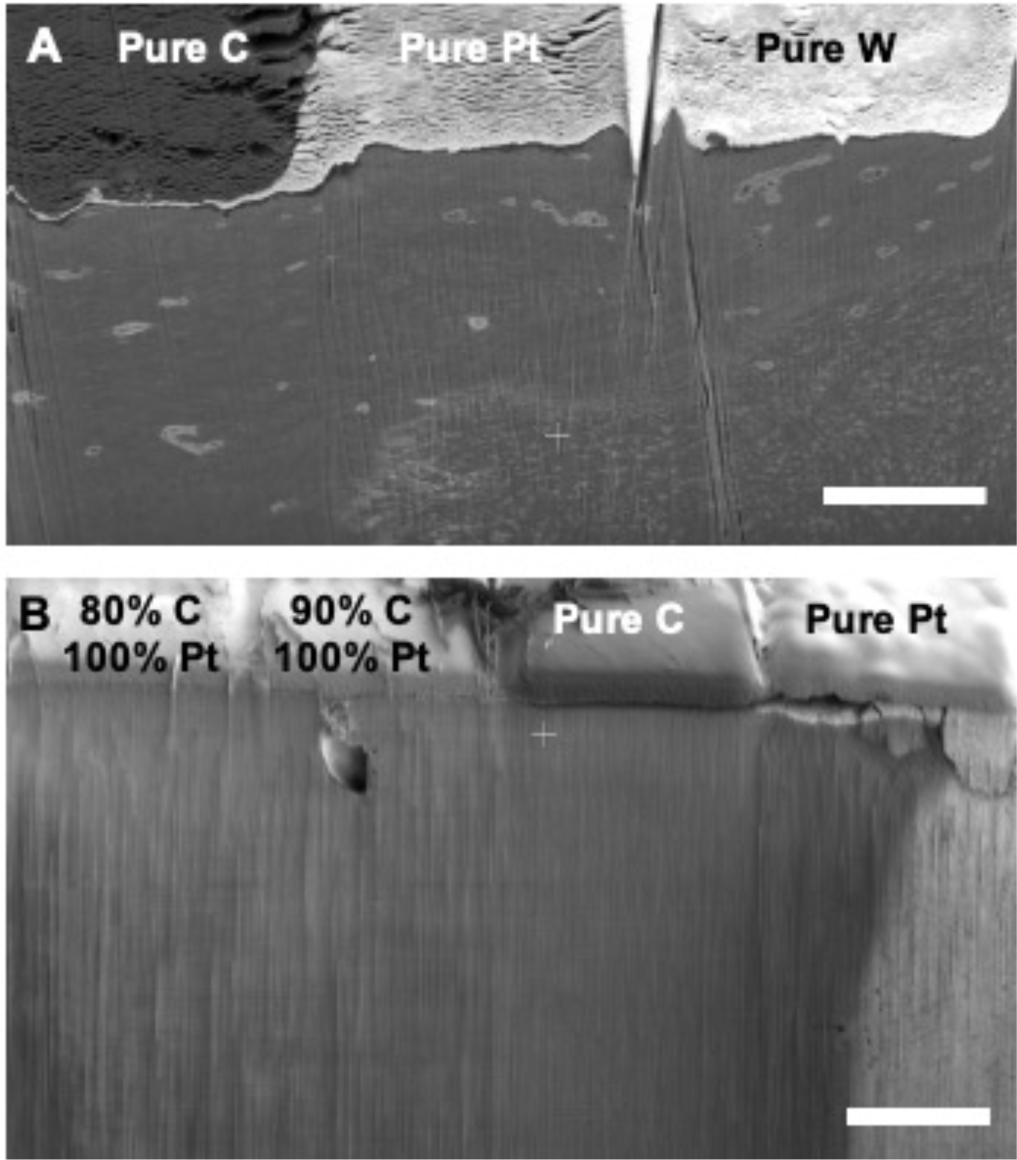
Capping layer composition experiments on demineralized and mineralized bone (A) Deposition of C, Pt, and W on demineralized bone after a small 30 slice tomography acquisition. A reduction in curtaining was noted under the Pt and W deposition layers. Scale bar: 5 μm. (B) Deposition of C, Pt, and C/Pt mixtures on mineralized bone after a 5 slice tomography acquisition. A clear reduction in curtaining was observed under the C deposition layer. Percentages refer to the flux rate of gas relative to the standard deposition flux rate in the PFIB. Scale bar: 20 μm.

**Figure S3.**
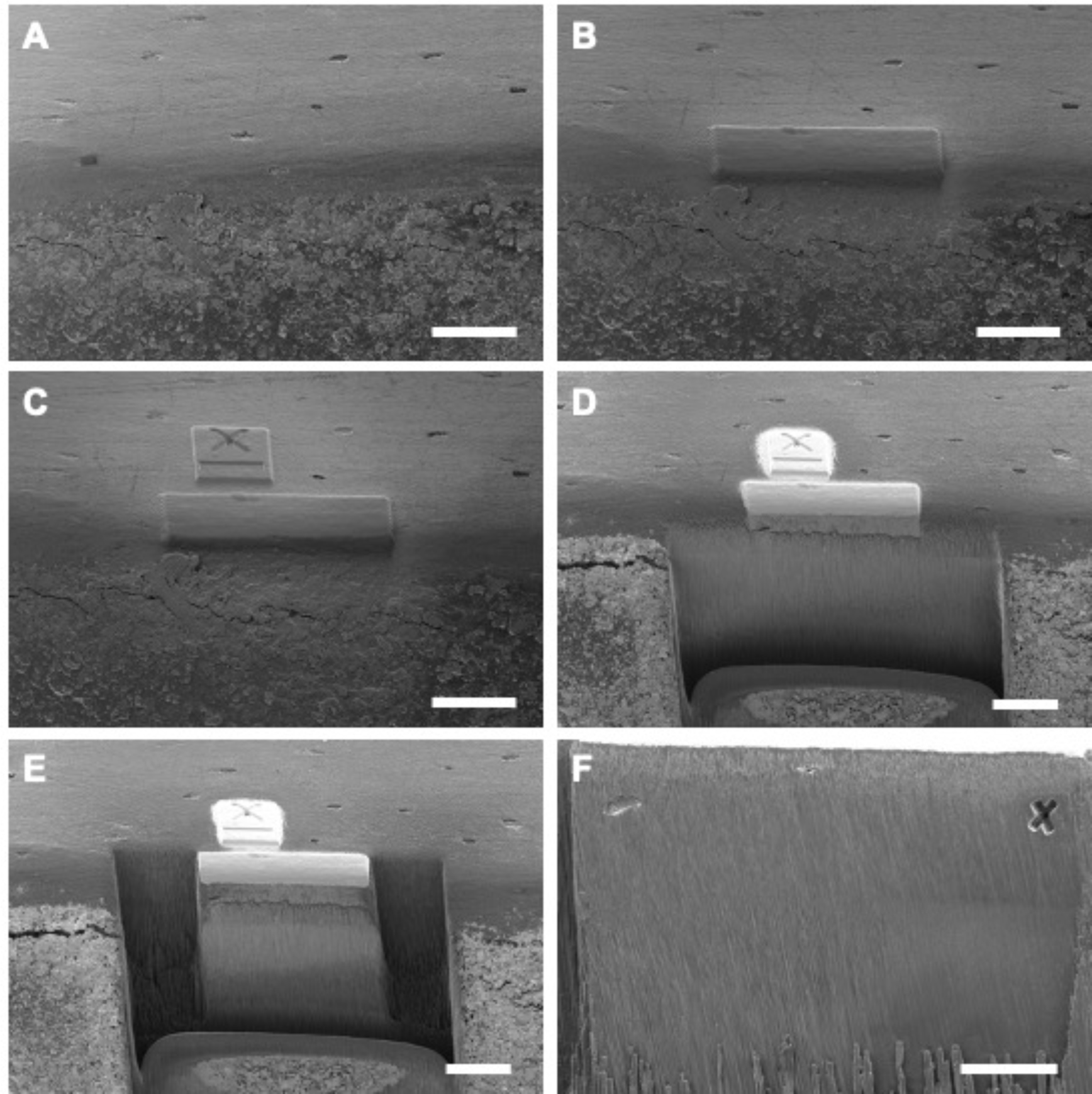
PFIB-SEM serial sectioning process. (A) A region of interest is selected. (B) The optimal protective capping layer is deposited on the region of interest, in this case, this is on the surface of the transverse plane. (C) A FIB fiducial is deposited and milled such that area can be tracked during automated serial sectioning. (D) A large cross section is exposed, this represents the XY milling and image plane. (E) Side trenches are milled to prevent shadowing and re-deposition. (F) A SEM fiducial is milled onto the cross-section to allow for easier image post-processing. All scale bars are 25 μm.

**Table S3.** Summary of FIB-SEM serial sectioning on mineralized tissues. Where not available, slice thickness (Z) is given in place of voxel size.

| Tissue | Preparation | Tomogram Size | Voxel Size | Reference |
| --- | --- | --- | --- | --- |
| **Human Bone** |  |  |  |  |
| Lamellar bone | Demineralized | 45.8 x 40.9 x 8.7 $\mu\text{m}$ | 20 nm | Present work |
| Lamellar bone | Mineralized | 28.6 x 25.6 x 8.7 $\mu\text{m}$ | 12.5 x 12.5 x 25 nm | Present work |
| Lamellar bone | Demineralized | N/A | 10 x 10 x 10 nm | Reznikov et al 2018 |
| Lamellar bone | Demineralized | Z = 6 – 9 $\mu\text{m}$ | 10 x 10 x 10 nm to 12.5 x 2.5 x 12.5 nm | Reznikov et al 2014 |
| Trabecular Bone | Demineralized | Z = 6 – 9 $\mu\text{m}$ | 10 x 10 x 10 nm to 12.5 x 12.5 x 12.5 nm | Reznikov et al 2015 |
| **Other Mammalian Bone** |  |  |  |  |
| Minipig Alveolar bone | Demineralized | Z = 10 – 15 $\mu\text{m}$ | 10 x 10 x 10 nm to 12.5 x 12.5 x 12.5 nm | Maria et al 2019 |
| Minipig fibrolamellar bone | Demineralized | X = 10.24 $\mu\text{m}$ Y= 8.6 $\mu\text{m}$ | 10 x 10 x 10 nm | Magal et al 2013 |
| Mouse trabecular bone | Mineralized | 25 x 20 x 20 $\mu\text{m}$ | Z = 50 nm | Hasegawa et al 2018 |
| Mouse proximal tibial metaphysis | Demineralized | 40 x 40 x 40 $\mu\text{m}$ | 20 x 20 x 20 nm | Robles et al 2019 |
| Mouse femur | Mineralized | 19.0 x 14 x 11 $\mu\text{m}$ | 18 x 18 x 29.5 nm | Schneider et al Bone 2011 |
| Mouse bone-cartridge interface | Mineralized | 30 x 20 x 17.5 $\mu\text{m}$ | Z = 50 nm | Haseqawa 2017 |
| Rat lamellar bone | Demineralized | 10 x 10 x 2 – 10 $\mu\text{m}$ | Z = 10 nm | Reznikov et al. Bone 2013 676 - 683 |
| **Avian Bone (Embryotic Chickens)** |  |  |  |  |
| Long bones | Cryo SEM | N/A | N/A | Kerschnitzki et al 2016 |
| Calvaria bone | Mineralized | 25 x 25 x 25 $\mu\text{m}$ | 25 x 25 x 25 nm | Hasimoto et al 2017 |
| **Tendon** |  |  |  |  |
| Turkey | Mineralized | N/A | 12 x 12 x 24 nm | Zou et al. 2019 |
| Turkey | Demineralized | N/A | 6 x 6 x 8.7 nm | Zou et al. 2019 |
| Rat | Demineralized | Z = 260 $\mu\text{m}$ | Z = 60 nm | Kanazawa et al 2014 |
| **Tooth Materials** |  |  |  |  |
| Mice cementum | Mineralized | 73 x 63 $\mu\text{m}$ x 8 $\mu\text{m}$ | 36 x 36 x 100 nm | Hirashima et al 2020 |
| Rat dentin | Demineralized | 73 x 63.6 x 4.2 $\mu\text{m}$ | 36 x 36 x 100 nm | Tanou et al 2018 |
| **Zebra fish** |  |  |  |  |
| Zebra fish | Demineralized | 26 x 22 x 12 $\mu\text{m}$ | 26 x 26 x 26 nm | Silvent et al 2017 |
| Zebra fish | Mineralized | 26 x 22 x 12 $\mu\text{m}$ | 26 x 26 x 26 nm | Silvent et al 2017 |
| Larval zebra fish | Mineralized, cryo | 30 x 23 x 12 $\mu\text{m}$ | 10 x 10 x 20 nm | Akiva et al 2019 |
| Larval zebra fish | Mineralized, cryo | 40 x 23 x 24.5 $\mu\text{m}$ | 20 x 20 x 40 nm | Akiva et al 2019 |

**Figure S4.**
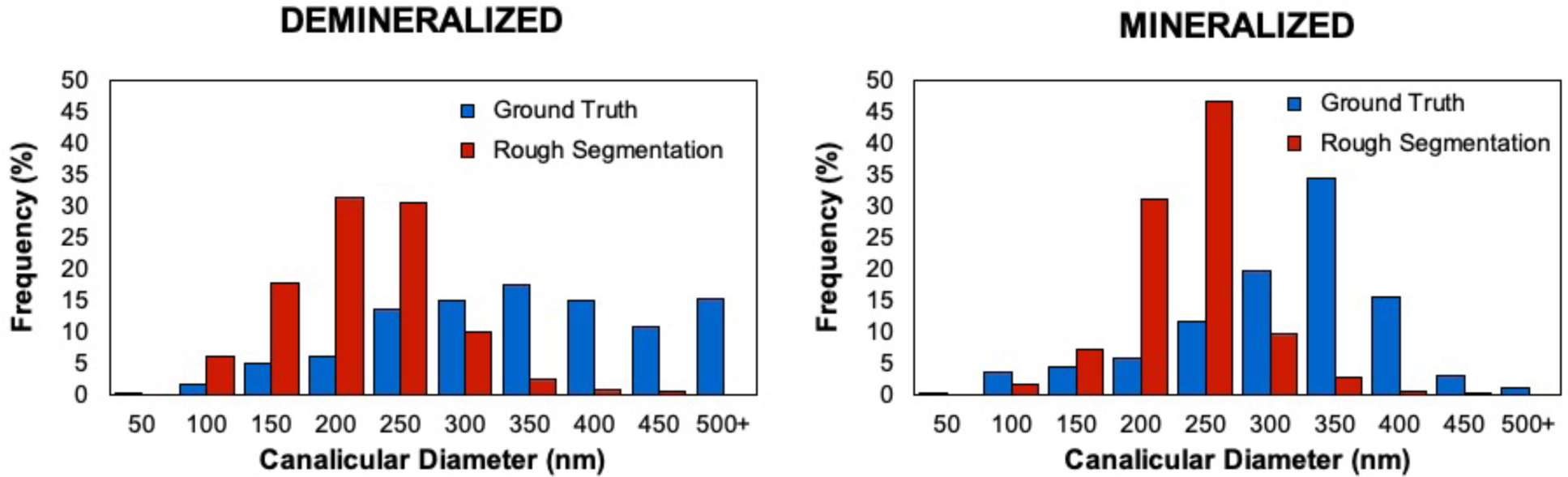
Dependence on neural network training quality. Results of two separate training inputs for (A) Demineralized tissue (B) Mineralized tissue. In the ground truth segmentations, training slices were diligently prepared where canalicular edges and interiors were included with only a small margin of error in guided segmentation. Rough segmentations included omissions of canaliculi or incomplete identification within single canaliculi, resulting in an apparent downward shift in the average canalicular diameter. A sufficient number of epochs were run in all cases to cause weighting factor convergence.

**Figure S5.**
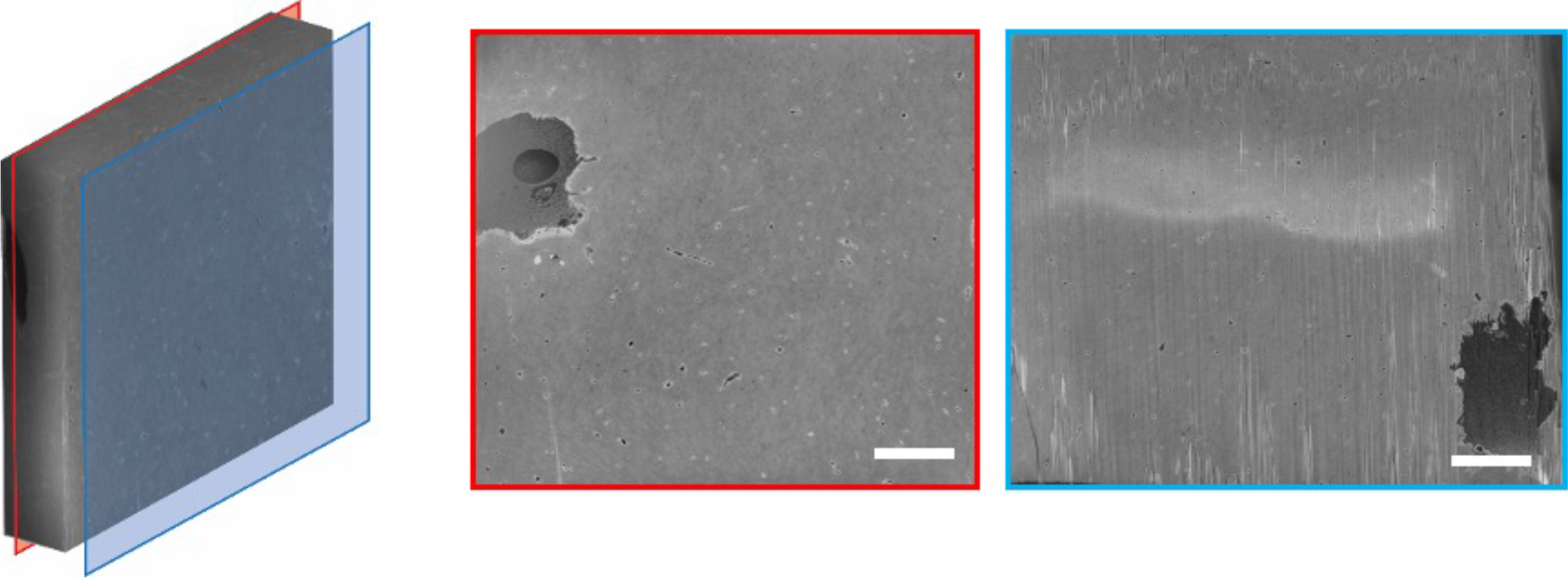
Two osteocytes present in demineralized bone. The demineralized dataset represents canalicular networks from two adjacent cells. The first osteocyte, bottom right of the blue plane, was encountered during preliminary milling. As PFIB serial sectioning requires a fine surface polish, several of the first slices, including this blue plane, are outside of the final volume recorded in the Slice and View tomogram. The second osteocyte, the top left of the red plane, is the osteocyte featured in the main manuscript. This juxtaposition of the two cells may account for the higher canalicular density observed in the demineralized dataset. Scale bar: 5 μm.

**Figure S6.**
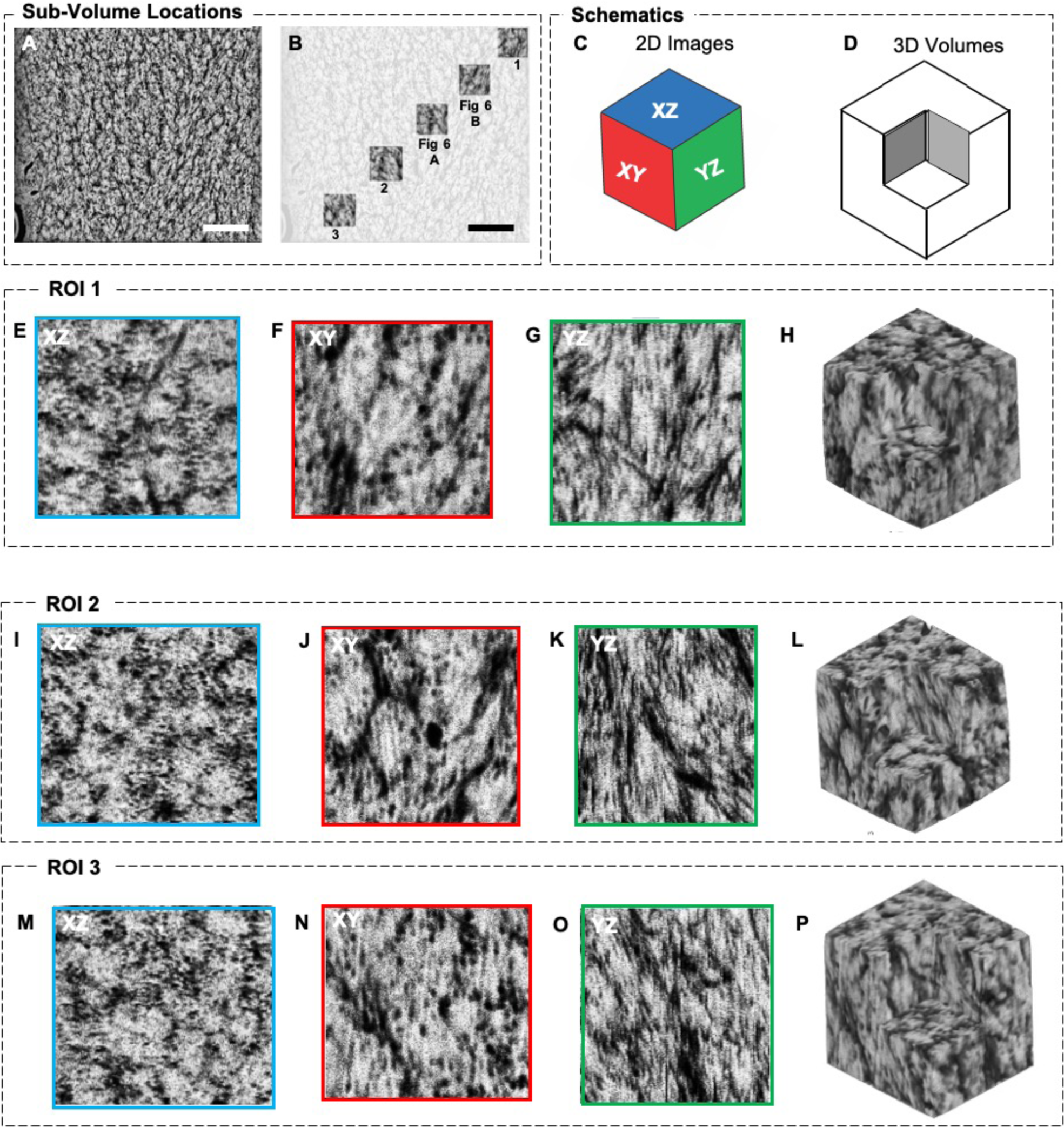
Exploring the mineral cluster orientation in 3D. Several additional ROIs (ROI 1,2,3) to the main manuscript Figures 6A and B are included here across the mineralized dataset. Consistent orientation of the mineralized features is noted, with rosettes appearing in the XZ (or transverse to the long axis of the femur), while collagen banding is regularly visible in the YZ plane along the length of the femur. The marquise-, or diamond-shaped motif, which makes up the ellipsoid in 3D space is consistently visible throughout the XY and YZ planes. ROIs measure 3 × 3×3 μm3. Scale bar: 1 μm.

## Movie Captions

**Movie S1. Orthogonal slices and LCN volume rendering of the demineralized dataset**. From 0 – 0:30 seconds, demineralized bone tomogram rotates 360°. 0:30 – 1:00 SEM images slice one-by-one in the XY plane from the front of the volume to the back. 1:00 – 1:30 SEM images slice one by one in the XY plane from the back of the volume to the front. 1:30 – 2:00 Tomogram disappears and the reconstructed LCN appear and rotate 360°. 2:00 – 2:30 Tomogram appears with the LCN and rotates 360°.

**Movie S2. Orthogonal slices and LCN volume rendering of the mineralized dataset**. From 0 – 0:30 seconds, mineralized bone tomogram rotates 360°. 0:30 – 1:00 SEM images slice one by one in the XY plane from the front of the volume to the back. 1:00 – 1:30 SEM images slice one by one in the XY plane from the back of the volume to the front. 1:30 – 2:00 Tomogram disappears and the reconstructed LCN appear and rotate 360°. 2:00 – 2:30 Tomogram appears with the LCN and rotates 360°.

**Movie S3. The morphology of the mineral clusters**. Orthogonal slices of marquise or ellipsoidal shapes in mineralized bone. An isolated mineral marquise (white) shown with orthogonal slices in the XY and XZ planes while rotating. At 10 seconds, the top-down view is shown, with the shape of a rosette apparent.

**Movie S4. 2D slice-by-slice of rosettes**. Single reconstructed slices from the XZ plane showing a zoomed in rosette shape as visualized in PFIB-SEM.

## Notes

### Competing Interest Statement

The authors have declared no competing interest.

## References

Addison, W.N., Nelea, V., Chicatun, F., Chien, Y.-C., Tran-Khanh, N., Buschmann, M.D., Nazhat, S.N., Kaartinen, M.T., Vali, H., Tecklenburg, M.M., Franceschi, R.T., McKee, M.D., 2014. Extracellular matrix mineralization in murine MC3T3-E1 osteoblast cultures: an ultrastructural, compositional and comparative analysis with mouse bone. Bone 71, 244–56. https://doi.org/10.1016/j.bone.2014.11.003

Akiva, A., Nelkenbaum, O., Schertel, A., Yaniv, K., Weiner, S., Addadi, L., 2019. Intercellular Pathways from the Vasculature to the Forming Bone in the Zebrafish Larval Caudal Fin: Possible Role in Bone Formation. J Struct Biol 206, 139–148. https://doi.org/10.1016/j.jsb.2019.02.011

Bassim, N., Scott, K., Giannuzzi, L.A., 2014. Recent advances in focused ion beam technology and applications. MRS Bulletin 39, 317–325. https://doi.org/10.1557/mrs.2014.52

Bernard, G.W., Pease, D.C., 1969. An electron microscopic study of initial intramembranous osteogenesis. Am J Anat 125, 271–290. https://doi.org/10.1002/aja.1001250303

Bigi, A., Ripamonti, A., Koch, M.H.J., Roveri, N., 1988. Calcified turkey leg tendon as structural model for bone mineralization. International Journal of Biological Macromolecules 10, 282–286–282–286.

Bonewald, L.F., 2011. The amazing osteocyte. J. Bone Miner. Res. 26, 229–238.

Bonucci, E., 1971. The Locus of Initial Calcification in Cartilage. Clin Orthop Relat R 78, 108–139. https://doi.org/10.1097/00003086-197107000-00010

Burnett, T.L., Kelley, R., Winiarski, B., Contreras, L., Daly, M., Gholinia, A., Burke, M.G., Withers, P.J., 2016. Large volume serial section tomography by Xe Plasma FIB dual beam microscopy. Ultramicroscopy 161, 119–129. https://doi.org/10.1016/j.ultramic.2015.11.001

Ciani, C., Doty, S.B., Fritton, S.P., 2009. An effective histological staining process to visualize bone interstitial fluid space using confocal microscopy. Bone 44, 1015–1017.

Currey, J.D., 2002. Bones: structure and mechanics. Princeton University Press.

Dong, P., Haupert, S., Hesse, B., Langer, M., Gouttenoire, P.-J., Bousson, V., Peyrin, F., 2014. 3D osteocyte lacunar morphometric properties and distributions in human femoral cortical bone using synchrotron radiation micro-CT images. Bone 60, 172–185.

Genthial, R., Beaurepaire, E., Schanne-Klein, M.-C., Peyrin, F., Farlay, D., Olivier, C., Bala, Y., Boivin, G., Vial, J.-C., Débarre, D., Gourrier, A., 2017. Label-free imaging of bone multiscale porosity and interfaces using third-harmonic generation microscopy. Scientific Reports 7, 3419.

Georgiadis, M., Müller, R., Schneider, P., 2016. Techniques to assess bone ultrastructure organization: orientation and arrangement of mineralized collagen fibrils. J Roy Soc Interface 13, 20160088. https://doi.org/10.1098/rsif.2016.0088

Giraud-Guille, M.M., 1988. Twisted plywood architecture of collagen fibrils in human compact bone osteons. Calcif. Tissue Int. 42, 167–180.

Grandfield, K., Vuong, V., Schwarcz, H.P., 2018. Ultrastructure of Bone: Hierarchical Features from Nanometer to Micrometer Scale Revealed in Focused Ion Beam Sections in the TEM. Calcified Tissue Int 103, 606–616. https://doi.org/10.1007/s00223-018-0454-9

Granke, M., Gourrier, A., Rupin, F., Raum, K., Peyrin, F., Burghammer, M., Saïed, A., Laugier, P., 2013. Microfibril Orientation Dominates the Microelastic Properties of Human Bone Tissue at the Lamellar Length Scale. Plos One 8, e58043. https://doi.org/10.1371/journal.pone.0058043

Hannah, K.M., Thomas, C.D.L., Clement, J.G., Carlo, F.D., Peele, A.G., 2010. Bimodal distribution of osteocyte lacunar size in the human femoral cortex as revealed by micro-CT. Bone 47, 866–871.

Hasegawa, T., Endo, T., Tsuchiya, E., Kudo, A., Shen, Z., Moritani, Y., Abe, M., Yamamoto, T., Hongo, H., Tsuboi, K., Yoshida, T., Nagai, T., Khadiza, N., Yokoyama, A., Freitas, P.H.L. de, Li, M., Amizuka, N., 2017. Biological application of focus ion beam-scanning electron microscopy (FIB-SEM) to the imaging of cartilaginous fibrils and osteoblastic cytoplasmic processes. J Oral Biosci 59, 55–62. https://doi.org/10.1016/j.job.2016.11.004

Hasegawa, T., Yamamoto, T., Hongo, H., Qiu, Z., Abe, M., Kanesaki, T., Tanaka, K., Endo, T., Freitas, P.H.L. de, Li, M., Amizuka, N., 2018. Three-dimensional ultrastructure of osteocytes assessed by focused ion beam-scanning electron microscopy (FIB-SEM). Histochem Cell Biol 149, 423–432. https://doi.org/10.1007/s00418-018-1645-1

Hirashima, S., Kanazawa, T., Ohta, K., Nakamura, K., 2020a. Three-dimensional ultrastructural imaging and quantitative analysis of the periodontal ligament. Anat Sci Int 95, 1–11. https://doi.org/10.1007/s12565-019-00502-5

Hirashima, S., Ohta, K., Kanazawa, T., Togo, A., Tsuneyoshi, R., Kusukawa, J., Nakamura, K., 2020b. Cellular network across cementum and periodontal ligament elucidated by FIB/SEM tomography. Microscopy 69, 53–58. https://doi.org/10.1093/jmicro/dfz117

Hu, C., Aindow, M., Wei, M., 2017. Focused ion beam sectioning studies of biomimetic hydroxyapatite coatings on Ti-6Al-4V substrates. Surf Coatings Technology 313, 255–262. https://doi.org/10.1016/j.surfcoat.2017.01.103

Kamioka, H., Kameo, Y., Imai, Y., Bakker, A.D., Bacabac, R.G., Yamada, N., Takaoka, A., Yamashiro, T., Adachi, T., Klein-Nulend, J., 2012. Microscale fluid flow analysis in a human osteocyte canaliculus using a realistic high-resolution image-based three-dimensional model. Integr Biol 4, 1198–1206. https://doi.org/10.1039/c2ib20092a

Kerschnitzki, M., Akiva, A., Shoham, A.B., Asscher, Y., Wagermaier, W., Fratzl, P., Addadi, L., Weiner, S., 2016a. Bone mineralization pathways during the rapid growth of embryonic chicken long bones. J Struct Biol 195, 82–92. https://doi.org/10.1016/j.jsb.2016.04.011

Kerschnitzki, M., Akiva, A., Shoham, A.B., Koifman, N., Shimoni, E., Rechav, K., Arraf, A.A., Schultheiss, T.M., Talmon, Y., Zelzer, E., Weiner, S., Addadi, L., 2016b. Transport of membrane-bound mineral particles in blood vessels during chicken embryonic bone development. Bone 83, 65–72. https://doi.org/10.1016/j.bone.2015.10.009

Kerschnitzki, M., Wagermaier, W., Roschger, P., Seto, J., Shahar, R., Duda, G.N., Mundlos, S., Fratzl, P., 2011. The organization of the osteocyte network mirrors the extracellular matrix orientation in bone. J. Struct. Biol. 173, 303–311.

Kollmannsberger, P., Kerschnitzki, M., Repp, F., Wagermaier, W., Weinkamer, R., Fratzl, P., 2017. The small world of osteocytes: connectomics of the lacuno-canalicular network in bone. New Journal of Physics 19, 073019–073019.

Landis, W.J., Hodgens, K.J., Arena, J., Song, M.J., McEwen, B.F., 1996. Structural relations between collagen and mineral in bone as determined by high voltage electron microscopic tomography. Microsc Res Techniq 33, 192–202.

Landis, W.J., Moradian-Oldak, J., Weiner, S., 1991. Topographic imaging of mineral and collagen in the calcifying Turkey tendon. Connective tissue research 25, 181–196.

Lee, B.E.J., Luo, L., Grandfield, K., Andrei, C.M., Schwarcz, H.P., 2019. Identification of collagen fibrils in cross sections of bone by electron energy loss spectroscopy (EELS). Micron 124, 102706. https://doi.org/10.1016/j.micron.2019.102706

Lin, Y., Xu, S., 2010. AFM analysis of the lacunar-canalicular network in demineralized compact bone. Journal of Microscopy 241, 291–302.

Loeber, T.H., Laegel, B., Wolff, S., Schuff, S., Balle, F., Beck, T., Eifler, D., Fitschen, J.H., Steidl, G., 2017. Reducing curtaining effects in FIB/SEM applications by a goniometer stage and an image processing method. J Vac Sci Technology B Nanotechnol Microelectron Mater Process Meas Phenom 35, 06GK01. https://doi.org/10.1116/1.4991638

Maria, R., Ben-Zvi, Y., Rechav, K., Klein, E., Shahar, R., Weiner, S., 2019. An unusual disordered alveolar bone material in the upper furcation region of minipig mandibles: a 3D hierarchical structural study. J Struct Biol 206, 128–137. https://doi.org/10.1016/j.jsb.2019.02.010

McNally, E.A., Schwarcz, H.P., Botton, G.A., Arsenault, A.L., 2012. A Model for the Ultrastructure of Bone Based on Electron Microscopy of Ion-Milled Sections. Plos One 7, e29258. https://doi.org/10.1371/journal.pone.0029258

Midura, R.J., Vasanji, A., Su, X., Midura, S.B., Gorski, J.P., 2008. Isolation of Calcospherulites from the Mineralization Front of Bone. Cells Tissues Organs 189, 75–79. https://doi.org/10.1159/000152914

Nudelman, F., Lausch, A.J., Sommerdijk, N.A.J.M., Sone, E.D., 2013. In vitro models of collagen biomineralization. J Struct Biol 183, 258–269. https://doi.org/10.1016/j.jsb.2013.04.003

Pacureanu, A., Langer, M., Boller, E., Tafforeau, P., Peyrin, F., 2012. Nanoscale imaging of the bone cell network with synchrotron X-ray tomography: optimization of acquisition setup. Med. Phys. 39, 2229.

Pacureanu, A., Larrue, A., Langer, M., Olivier, C., Muller, C., Lafage-Proust, M.-H., Peyrin, F., 2013. Adaptive filtering for enhancement of the osteocyte cell network in 3D microtomography images. IRBM 34, 48–52. https://doi.org/10.1016/j.irbm.2012.12.013

Peyrin, F., Dong, P., Pacureanu, A., Langer, M., 2014. Micro- and Nano-CT for the Study of Bone Ultrastructure. Curr Osteoporos Rep 12, 465–474. https://doi.org/10.1007/s11914-014-0233-0

Peyrin, F., Pacureanu, A., Zuluaga, M.A., Dong, P., Langer, M., 2012. 3D X-Ray CT imaging of the bone lacuno-canalicular network. 2012 9th IEEE Intl Symposium Biomedical Imaging (ISBI) 1, 1788–1791. https://doi.org/10.1109/isbi.2012.6235929

Repp, F., Kollmannsberger, P., Roschger, A., Kerschnitzki, M., Berzlanovich, A., Gruber, G.M., Roschger, P., Wagermaier, W., Weinkamer, R., 2017. Spatial heterogeneity in the canalicular density of the osteocyte network in human osteons. Bone Reports 6, 101–108.

Reznikov, N., Almany-Magal, R., Shahar, R., Weiner, S., 2013. Three-dimensional imaging of collagen fibril organization in rat circumferential lamellar bone using a dual beam electron microscope reveals ordered and disordered sub-lamellar structures. Bone 52, 676–683. https://doi.org/10.1016/j.bone.2012.10.034

Reznikov, N., Bilton, M., Lari, L., Stevens, M.M., Kröger, R., 2018. Fractal-like hierarchical organization of bone begins at the nanoscale. Science 360, eaao2189. https://doi.org/10.1126/science.aao2189

Reznikov, N., Chase, H., Brumfeld, V., Shahar, R., Weiner, S., 2014a. The 3D structure of the collagen fibril network in human trabecular bone: relation to trabecular organization. Bone 71, 189–95. https://doi.org/10.1016/j.bone.2014.10.017

Reznikov, N., Shahar, R., Weiner, S., 2014b. Three-dimensional structure of human lamellar bone: The presence of two different materials and new insights into the hierarchical organization. Bone 59, 93–104. https://doi.org/10.1016/j.bone.2013.10.023

Reznikov, N., Shahar, R., Weiner, S., 2014c. Bone hierarchical structure in three dimensions. Acta Biomater 10, 3815–3826. https://doi.org/10.1016/j.actbio.2014.05.024

Sano, H., Kikuta, J., Furuya, M., Kondo, N., Endo, N., Ishii, M., 2015. Intravital bone imaging by two-photon excitation microscopy to identify osteocytic osteolysis in vivo. Bone 74, 134–139.

Schneider, P., Meier, M., Wepf, R., Müller, R., 2011. Serial FIB/SEM imaging for quantitative 3D assessment of the osteocyte lacuno-canalicular network. Bone 49, 304–11. https://doi.org/10.1016/j.bone.2011.04.005

Schwarcz, H.P., 2015. The ultrastructure of bone as revealed in electron microscopy of ion-milled sections. Semin Cell Dev Biol 46, 44–50. https://doi.org/10.1016/j.semcdb.2015.06.008

Shah, F.A., Palmquist, A., 2017. Evidence that Osteocytes in Autogenous Bone Fragments can Repair Disrupted Canalicular Networks and Connect with Osteocytes in de novo Formed Bone on the Fragment Surface. Calcified Tissue Int 101, 321–327. https://doi.org/10.1007/s00223-017-0283-2

Shah, F.A., Ruscsák, K., Palmquist, A., 2020. Transformation of bone mineral morphology: From discrete marquise-shaped motifs to a continuous interwoven mesh. Bone Reports 100283. https://doi.org/10.1016/j.bonr.2020.100283

Shah, F.A., Zanghellini, E., Matic, A., Thomsen, P., Palmquist, A., 2016. The Orientation of Nanoscale Apatite Platelets in Relation to Osteoblastic–Osteocyte Lacunae on Trabecular Bone Surface. Calcified Tissue Int 98, 193–205. https://doi.org/10.1007/s00223-015-0072-8

Silvent, J., Akiva, A., Brumfeld, V., Reznikov, N., Rechav, K., Yaniv, K., Addadi, L., Weiner, S., 2017. Zebrafish skeleton development: High resolution micro-CT and FIB-SEM block surface serial imaging for phenotype identification. Plos One 12, e0177731. https://doi.org/10.1371/journal.pone.0177731

Varga, P., Hesse, B., Langer, M., Schrof, S., Männicke, N., Suhonen, H., Pacureanu, A., Pahr, D., Peyrin, F., Raum, K., 2015. Synchrotron X-ray phase nano-tomography-based analysis of the lacunar-canalicular network morphology and its relation to the strains experienced by osteocytes in situ as predicted by case-specific finite element analysis. Biomechanics and Modeling in Mechanobiology 14, 267–282.

Varga, P., Weber, L., Hesse, B., Langer, M., 2016. X-ray and Neutron Techniques for Nanomaterials Characterization 1–42. https://doi.org/10.1007/978-3-662-48606-1_1

Wagermaier, W., Gupta, H.S., Gourrier, A., Burghammer, M., Roschger, P., Fratzl, P., 2006. Spiral twisting of fiber orientation inside bone lamellae. Biointerphases 1, 1–5.

Wang, X., Shah, F.A., Palmquist, A., Grandfield, K., 2016. 3D Characterization of Human Nano-osseointegration by On-Axis Electron Tomography without the Missing Wedge. Acs Biomater Sci Eng 3, 49–55. https://doi.org/10.1021/acsbiomaterials.6b00519

Weiner, S., Traub, W., 1992. Bone structure: from angstroms to microns. FASEB 6, 879–885. https://doi.org/10.1096/fasebj.6.3.1740237

Weiner, S., Traub, W., 1989. Crystal size and organization in bone. Connect Tissue Res 21, 259–65. https://doi.org/10.3109/03008208909050015

Weiner, S., Traub, W., Wagner, H.D., 1999. Lamellar bone: structure-function relations. J. Struct. Biol. 126, 241–255.

Wittig, N.K., Laugesen, M., Birkbak, M.E., Bach-Gansmo, F.L., Pacureanu, A., Bruns, S., Wendelboe, M.H., Brüel, A., Sørensen, H.O., Thomsen, J.S., Birkedal, H., 2019. Canalicular Junctions in the Osteocyte Lacuno-Canalicular Network of Cortical Bone. ACS Nano 13, 6421–6430. https://doi.org/10.1021/acsnano.8b08478

You, L., Weinbaum, S., Cowin, S.C., Schaffler, M.B., 2004. Ultrastructure of the osteocyte process and its pericellular matrix. Anatomical Rec Part Discov Mol Cell Evol Biology 278A, 505–513. https://doi.org/10.1002/ar.a.20050

Zou, Z., Tang, T., Macías-Sánchez, E., Sviben, S., Landis, W.J., Bertinetti, L., Fratzl, P., 2019. Three-dimensional Structural Interrelations between Cells, Extracellular Matrix and Mineral in Vertebrate Mineralization. BioRxiv 803007. https://doi.org/10.1101/803007

## Supplementary References

Utke,I., Hoffmann,P., Melngailis, J., 2008. Gas-assisted focused electron beam and ion beam processing and fabrication. J Vac Sci Technology B Microelectron Nanometer Struct 26, 1197.https://doi.org/10.1116/1.2955728

